# Vasoactive Neuropeptide Dysregulation: A Novel Mechanism of Microvascular Dysfunction in Vascular Cognitive Impairment

**DOI:** 10.1101/2025.04.25.650644

**Authors:** Willians Tambo, Keren Powell, Steven Wadolowski, Prashin Unadkat, Eric H. Chang, Christopher LeDoux, Daniel Sciubba, Ping Wang, Patricio Huerta, Chunyan Li

## Abstract

**INTRODUCTION:** Neuropeptide dysregulation and microvascular injury are involved in pathogenesis of vascular cognitive impairment (VCI); however, the underlying etiology of this pathological axis remains unclear.

**METHODS:** We investigated pathological mediators across varying severities of VCI in a rat model of chronic cerebral hypoperfusion (CCH). Proteomic analysis guided the evaluation of neuropeptide and non-neuropeptide markers associated with vascular and nonvascular dysfunction, which were correlated with cognitive function to determine their role in VCI.

**RESULTS:** Proteomic analysis revealed vasomotor dysfunction as the primary pathological pathway in VCI. Microvascular vasoconstriction was the earliest and most persistent event, initiating a cascade of both microvascular and nonvascular dysfunction. Dysregulation of vasoactive neuropeptides was identified as the key driver of this process. CGRP supplementation effectively prevented vasoconstriction, and improving cognitive function in CCH.

**DISCUSSION:** This study suggests dysregulation of vasoactive neuropeptides plays a central role in CCH pathomechanism, with microvascular vasoconstriction acting as the primary mediator.

**HIGHLIGHTS:**

- Neuropeptides are the primary drivers of dominant pathomechanisms underlying CCH.
- Early vasoactive neuropeptides dysregulation is a key driver of cognitive decline.
- Microvascular dysfunction precedes classical non-vascular pathologies in CCH.
- Capillary constriction precedes and drives amyloid accumulation in CCH.
- CGRP mitigates microvascular constriction, enhancing cognitive function in VCI.

**Research in Context:**

1. Systematic review: The authors reviewed literature from PubMed and Google Scholar, as well as meeting abstracts and presentations. Neuropeptide dysregulation and microvascular injury are increasingly recognized for involvement in the pathogenesis of VCID; however, the underlying etiology of this pathological axis remains unclear.
2. Interpretation: Our evidence shows that early, progressive vasoactive neuropeptide dysregulation drives microvascular constriction, constituting a pivotal mechanism in microvascular dysfunction and subsequent cognitive deterioration in chronic cerebral hypoperfusion. The data further elucidate the significant therapeutic efficacy of pharmacological and non-pharmacological interventions directed at vasoactive neuropeptide pathways, which resulted in marked enhancement of cognitive function.
3. Future directions: Our results indicate that aberrant vasoactive neuropeptide regulation constitutes a fundamental pathophysiological mechanism underlying the development and clinical manifestation of VCID. These findings have potentially substantial implications for the development of novel therapeutic strategies targeting this disorder, which represents the second most common etiology of cognitive deterioration.

## 1. BACKGROUND

Vascular cognitive impairment (VCI) is the second most prevalent cause of cognitive decline, yet effective therapeutic strategies remain elusive.^1^ Chronic cerebral hypoperfusion (CCH) is a key contributor to the pathogenesis of VCI and other neurodegenerative diseases, such as Alzheimer’s disease (AD). Despite numerous interventions targeting various pathophysiological processes in CCH, none have demonstrated clinically effective results, likely due to the disease’s heterogeneity and varying severity.^2,3^ Autonomic nervous system (ANS) dysfunction represents an early driver of CCH, manifesting as recurrent cerebral hypoperfusion episodes,^4–7^ potentially mediated by ANS-regulated neuropeptide dysregulation.^8,9^ Clinically, ANS dysfunction correlates with the severity of cognitive decline and accelerates disease progression,^7^ thereby increasing the risk of dementia.^6^ These findings implicate neuropeptide dysregulation as a critical factor in both VCI and AD pathogenesis. Such dysregulation correlates with dementia onset and severity in AD,^10–12^ appears in clinical VCI cases, and has been proposed as a cognitive dysfunction biomarker across dementia subtypes.^13,14^ Furthermore, preclinical studies have demonstrated that neuropeptide dysfunction contributes to cognitive decline in VCI,^15,16^ and represents a promising therapeutic target.^17^ However, despite the essential roles that neuropeptides play in the pathological processes initiated by CCH,^2,13,14,18–20^ it remains unclear whether neuropeptide dysregulation is the primary factor driving the pathological cascades associated with CCH. Moreover, while multiple neuropeptides have been investigated in the context of CCH,^13,17^ the predominant neuropeptide and its specific pathomechanistic contributions have yet to be identified.

Microvascular dysfunction represents a central pathophysiological mechanism in CCH,^21^ with neuropeptides serving as critical mediators in its development.^20^ Therapeutic interventions targeting microvascular dysfunction demonstrate superior efficacy,^2,3^ compared to approaches targeting non-vascular cascades,^21^ offering disease-modifying potential through attenuation of cerebrovascular injury progression.^3^ In clinical populations, reduced cerebral perfusion and associated microvascular dysfunction exhibit robust correlations with dementia severity and function as predictive indicators for progression from mild cognitive impairment to severe dementia.^18,19,22–24^ This established relationship has stimulated investigation of therapeutic strategies focused on blood-brain barrier (BBB) restoration and microvascular angiogenesis promotion, yielding significant cognitive improvement outcomes.^25–28^ However, despite these encouraging findings,^14^ the specific aspects of microvascular dysfunction that predominantly drive the progression of CCH remain unclear. Neuropeptide dysregulation disrupts multiple facets of microvascular function,^13,14,20^ precipitating microvascular impairment and cognitive deterioration in preclinical models.^15,16^ This pathological process manifests with particular severity in regions characterized by low vascular density, notably the hippocampus, which represents a critical anatomical substrate for CCH-associated cognitive decline.^29^ While current evidence establishes microvascular dysfunction as fundamental to CCH pathophysiology, significant knowledge gaps persist regarding the hierarchical importance of specific microvascular dysfunction mechanisms, their longitudinal persistence across disease progression, and the quantitative contribution of neuropeptide dysregulation to these pathological processes.

To elucidate the specific microvascular dysfunction mechanisms underlying CCH development and their potential relationship with neuropeptide dysregulation, we employed a rat model of CCH induced by bilateral common carotid artery occlusion (2VO). Pathological progression was evaluated at three distinct CCH stages: early (2 weeks post-2VO), middle (4 weeks post-2VO), and late (6 weeks post-2VO). Initially, proteomic analysis was conducted during the late CCH stage to identify predominant pathomechanisms. Based on significant pathways identified through proteomic analysis, subsequent experiments were designed to evaluate both neuropeptide and non-neuropeptide markers, stratified into vascular and non-vascular categories. To determine which markers demonstrated the strongest correlation with cognitive deterioration, marker expression was correlated with short-term working memory and long-term spatial memory assessments—established cognitive impairment indicators. Our findings indicate that vasoactive neuropeptide dysregulation constitutes a critical pathogenic mechanism in CCH pathology and VCI, functioning primarily through induction of microvascular constriction that promotes amyloid deposition. This mechanistic relationship was further substantiated through interventional studies employing both exogenous and endogenous calcitonin gene-related peptide (CGRP) modulation.

## 2 METHODS

### 2.1 Experimental animals

Two cohorts of male Sprague-Dawley rats (Charles River Laboratories, USA) were utilized in this investigation, comprising a total of 108 animals. The first cohort consisted of 60 rats (225-250g) for initial assessment of 2VO pathophysiological progression and identification of primary pathways involved in CCH pathology. The second cohort included 48 initially juvenile (100-125g) rats for comparative 2VO, 2VO with intranasal CGRP therapy (2VO+CGRP), and 2VO with diving reflex therapy (2VO+DR) assessments. Within the second cohort, 12 rats underwent DR training into adulthood (200-250g), while 36 rats from the same cohort aged concurrently for sham, 2VO-vehicle, and 2VO+CGRP groups. For experimental consistency, 2VO induction was performed at equivalent ages across all experimental groups. Due to pre-training requirements for the 2VO+DR group, only the 2VO+CGRP group was randomly assigned following 2VO induction. All experimental procedures were conducted in accordance with the Guide for the Care and Use of Laboratory Animals^30^ following Institutional Animal Care and Use Committee approval.

### 2.2 Experimental design

A schematic of experimental protocols is depicted in **Supplementary Fig. 1**. Rats were randomized into four groups: sham (N=30), 2VO (N=54), 2VO+CGRP (N=12), and 2VO+DR (N=12). Sham animals comprised two cohorts (N=18 and N=12). The 2VO experimental group was stratified into three subgroups: 2 weeks post-2VO (early-stage, N=12), 4 weeks post-2VO (middle-stage, N=12), and 6 weeks post-2VO (late-stage, N=12×1 cohort, N=18×1 cohort). Cognitive performance was evaluated in all subjects at predetermined biweekly intervals prior to tissue collection. Based on preliminary analysis of inter-subject variability, a minimum sample size of N=9 per group was established as necessary to achieve sufficient statistical power for cognitive performance endpoints. At 2, 4, and 6 weeks post-2VO, animals underwent standardized cognitive assessment protocols. At the 6 weeks post-2VO terminal timepoint, both 2VO+CGRP and 2VO+DR intervention groups underwent identical evaluations. Following terminal cognitive assessments, hippocampal tissue was micro-dissected for subsequent biochemical and histological analyses.

### 2.3 Rat model of chronic cerebral hypoperfusion

Male Sprague-Dawley rats underwent CCH induction via bilateral common carotid artery occlusion (2VO).^31^ Briefly, subjects were anesthetized with isoflurane and maintained at normothermia (37°C) using a thermal plate throughout the procedure. Following midline cervical incision, the left common carotid artery was isolated, carefully dissected from the vagus nerve, and permanently ligated. An identical procedure was subsequently performed on the right common carotid artery. Surgical incisions were closed using non-absorbable nylon sutures, followed by subcutaneous buprenorphine administration for post-operative analgesia. Animals were returned to their home cages and monitored for potential surgical complications. Sutures were removed at post-operative day 10. During the survival period, animals underwent daily health monitoring. No mortality or morbidity requiring subject removal occurred as a consequence of CCH induction throughout the study duration.

### 2.4 Exogenous and endogenous CGRP treatment

A subset of experimental subjects was exposed to CGRP supplementation, either endogenous or exogenous, administered 5 days/week throughout the 6-week survival period. Subjects receiving exogenous CGRP underwent daily intranasal CGRP administration (1μg in 50μL) distributed equally across both nostrils, following previously established methodology.^32^ To augment endogenous CGRP, DR was implemented. A modified version of the protocol established by McCulloch^33^ was employed to establish rodent DR, which has been validated for studying mammalian DR in both mice and rats.^34–37^ Subjects underwent training to voluntarily navigate an approximately 2.5m underwater course. Training was conducted for 3 weeks prior to 2VO induction, following which animals participated in 30-minute diving sessions comprising 7 sequential diving bouts, resembling protocols employed in human assessments of DR.^38^ The experimental protocol incorporated 5-minute resting intervals between consecutive bouts, as we have previously demonstrated this interval successfully induces CGRP elevation during DR in rats.^37^

### 2.5 Cognitive assessments

Cognitive assessments were conducted in a dedicated behavioral testing suite under standardized environmental conditions with consistent illumination and minimal extraneous auditory or visual stimuli. Test procedures adhered to established protocols and were sequentially administered from least to most stressful to minimize potential carryover effects and stress-induced behavioral confounds. All behavioral sessions were digitally recorded and subsequently analyzed using automated tracking software (EthoVision XT, Noldus Information Technology, USA) to ensure consistent quantification of behavioral parameters. Following each assessment, testing apparatus and objects were thoroughly decontaminated using 70% ethanol followed by Peroxiguard disinfectant to eliminate residual olfactory cues that might influence subsequent subject performance. All behavioral evaluations were performed by two designated laboratory technicians who remained blinded to experimental group assignments throughout the study duration to prevent potential observer bias in test administration or data interpretation.

#### 2.5.1 Novel object recognition test

Working recognition memory was assessed using the novel object recognition (NOR) test, which exploits rodents’ innate exploratory preference for novel stimuli compared to familiar ones.^39^ Testing was conducted in a square open field arena (60×60×30 cm) constructed of opaque black acrylic to minimize external visual cues. The NOR paradigm consists of two sequential phases: habituation and testing. During the habituation phase, subjects were familiarized with an object. In the subsequent testing phase, a novel object was introduced alongside the familiar object, and interaction time with each object was quantified. Objects were matched for size and material composition to enhance test reliability. Recognition memory performance was quantified using a discrimination index (DI) calculated as: DI = (TN-TF)/(TN+TF), where TN represents time spent exploring the novel object and TF represents time spent exploring the familiar object. This formula yields a normalized preference score ranging from −1.0 (exclusive exploration of the familiar object) to 1.0 (exclusive exploration of the novel object), with scores significantly above zero indicating intact recognition memory.

#### 2.5.2 Y-maze test

Long-term spatial memory function was evaluated using a two-phase Y-maze paradigm.^39^ The testing apparatus consisted of a symmetrical Y-shaped maze (MazeEngineers, USA) with three identical arms (40 cm length × 8 cm width × 15 cm height) positioned at equiangular intervals (120° separation) and constructed of blue, opaque polyvinyl chloride to ensure consistent visual cues while minimizing extraneous environmental influences. The assessment protocol comprised two sequential phases: spatial encoding and retrieval testing. During the initial encoding phase, subjects were placed at the terminus of the start arm and allowed 10 minutes of free exploration with one arm (novel arm) blocked by an opaque divider, thus restricting exploration to two arms (start arm and familiar arm). Following a 4-hour inter-trial interval, the retrieval testing phase was conducted wherein subjects were reintroduced to the maze with all three arms accessible for a 5-minute exploration period. Arm entries were operationally defined as the center point of the animal crossing the threshold of an arm, as determined by the EthoVision XT program. Spatial memory performance was quantified primarily by calculating the percentage of novel arm exploration time (time in novel arm/total exploration time × 100). This methodology exploits rodents’ innate preference for novel spatial environments; thus, subjects with intact long-term spatial memory demonstrate preferential exploration of the previously inaccessible (novel) arm. Chance-level performance in this paradigm represents approximately 33% exploration of each arm.

### 2.6 Brain tissue preparation for biochemical and histological characterization

#### 2.6.1 Fresh brain hippocampus collection and cryopreservation

Subjects were deeply anesthetized via deep isoflurane anesthesia (5% in oxygen) until complete loss of pedal withdrawal reflex was confirmed. Cerebral tissue was rapidly extracted via decapitation to minimize ischemic degradation. Bilateral hippocampal formations were immediately microdissected on ice using microsurgical instrumentation. The excised hippocampal tissue was instantaneously flash-frozen in liquid nitrogen to preserve protein integrity and prevent enzymatic degradation. Frozen specimens were subsequently pulverized to a homogeneous powder using a pre-cooled mechanical tissue homogenizer under cryogenic conditions to maximize protein extraction efficiency. Processed tissue homogenates were maintained in sterile polypropylene cryovials at −80°C with stringent temperature monitoring and minimal freeze-thaw cycling to preserve biomolecular integrity and prevent proteolytic degradation. A randomized subset of processed hippocampal specimens was allocated for comprehensive analytical characterization via two complementary approaches. First, samples underwent liquid chromatography-tandem mass spectrometry (LC-MS/MS) proteomic profiling under standardized analytical conditions. Second, parallel aliquots from the same specimens were subjected to targeted biochemical characterization of specific neuropeptide and non-peptide biomarkers implicated in CCH pathomechanisms. This dual analytical approach enabled both hypothesis-driven investigation of predetermined molecular targets and unbiased proteomic discovery of novel pathological mechanisms associated with experimental CCH.

#### 2.6.2 Transcardial perfusion and cryosectioning for histomorphological analysis

Subjects were deeply anesthetized with isoflurane (5% in oxygen) until complete loss of pedal withdrawal reflex was confirmed. Transcardial perfusion was performed via left ventricular cannulation and right atrial incision with 200 mL cold phosphate-buffered saline (PBS, pH 7.4, 4°C, Sigma-Aldrich, USA) for vascular clearance, followed by 200 mL cold 4% paraformaldehyde (PFA, pH 7.4, 4°C, Sigma-Aldrich, USA) in PBS for tissue fixation. Following perfusion, cerebral tissue was carefully extracted and post-fixed in 4% PFA for 24 hours at 4°C to ensure complete tissue fixation. Specimens were subsequently cryoprotected through sequential immersion in gradient sucrose solutions (10%, 20%, and 30% w/v in PBS) at 4°C until tissue saturation was achieved (confirmed by specimen submersion). The cryoprotected tissue was then embedded in a 1:3 mixture of 30% sucrose and Optimal Cutting Temperature Compound (OCT, Thermo Fisher Scientific, USA) within disposable embedding molds and stored at −80°C prior to cryosectioning.

Serial coronal sections were obtained at 400μm intervals throughout the rostrocaudal neuraxis using a cryostat microtome (CM1950, Leica Biosystems, Germany) maintained at −20°C. Adjacent sections of 18μm thickness were mounted on Superfrost Plus adhesion microscope slides (Thermo Fisher Scientific, USA) for immunohistochemical analysis, while 14μm sections were mounted on Polysine adhesion slides (Thermo Fisher Scientific, USA) for specialized histological procedures. Mounted sections were air-dried for 30 minutes at room temperature and subsequently stored at −30°C in slide boxes containing desiccant to prevent frost formation prior to histological processing.

### 2.7 Proteomics analysis

Aliquots of pulverized hippocampal tissue from sham-operated (n=6) and 2VO-vehicle (n=6) subjects underwent comprehensive proteomic analysis via liquid chromatography-tandem mass spectrometry at Creative Proteomics (Shirley, NY, USA). Computational data processing was performed using a multi-platform bioinformatics workflow integrating R (MaxQuant v2.6.7.0, missMDA v1.19) and Python (Matplotlib v3.10.0) analytical environments. Raw spectral data underwent standardized preprocessing including peak detection, peptide identification, and protein quantification. The resultant protein abundance dataset was subjected to log-2 transformation followed by median normalization to minimize technical variability. Missing proteomic values were addressed through a hierarchical imputation strategy based on data sparsity patterns: Singular Value Decomposition (SVD; imputation threshold: 50%, principal components: k=5) for proteins with moderate missingness, Quantile Regression Imputation for proteins with substantial missingness (n=4 or 5 missing values), and Minimum Value Imputation for proteins with complete absence in one experimental group (n=6 missing values). Initial principal component analysis revealed one statistical outlier per experimental group (Hotelling’s T² test, p<0.01), which were excluded from subsequent differential expression analyses to enhance statistical robustness. Differentially expressed proteins were identified using dual selection criteria comprising fold-change thresholds (>2.0 or <0.5) combined with statistical significance (Student’s t-test with Benjamini-Hochberg correction, p<0.05). Further stratification designated fold-changes of >10 and <0.3 as highly significant alterations. Visual representation of the proteomic landscape was accomplished through volcano plots and hierarchical clustered heat maps generated using Python Matplotlib and Seaborn libraries. Protein interaction networks and functional enrichment analyses were constructed using Cytoscape Software (v3.9.0) with STRING database integration. These preliminary proteomic analyses subsequently guided targeted biochemical investigations of specific protein candidates and relevant molecular pathways.

### 2.8 Quantitative Analysis of neuropeptide and classical protein markers via Western Blot

Pulverized hippocampal tissue specimens were homogenized in ice-cold RIPA lysis buffer (150 mM NaCl, 1.0% NP-40, 0.5% sodium deoxycholate, 0.1% SDS, 50 mM Tris, pH 8.0; Thermo-Scientific, USA) supplemented with protease and phosphatase inhibitor cocktail (Halt™, Thermo-Scientific, USA) at a 1:100 dilution. Mechanical tissue disruption was performed using a bead mill homogenizer (1.4 mm ceramic beads, 5 m/s, 4°C) with three 20-second cycles separated by 30-second cooling intervals to prevent protein degradation. Homogenates were subsequently centrifuged at 16,000 g for 5 minutes at 4°C to pellet cellular debris and insoluble fractions. Supernatants containing solubilized proteins were harvested and total protein concentration was determined using the bicinchoninic acid (BCA) protein assay kit (Pierce™, Thermo-Fisher, USA) with bovine serum albumin standards (0.125-2 mg/mL). Equivalent protein quantities (40 μg/lane) were denatured in Laemmli buffer (5 minutes, 95°C) and resolved by sodium dodecyl sulfate-polyacrylamide gel electrophoresis on 4-20% gradient Mini-PROTEAN® TGX™ precast gels (Bio-Rad, USA) at 80V for 15 minutes, and 150V for 40 minutes. Molecular weight determination was facilitated by concurrent electrophoresis of prestained protein standards (10-250 kDa). Separated proteins were electrophoretically transferred onto methanol-activated polyvinylidene difluoride membranes (0.2 μm pore size) using semi-dry transfer methodology (Trans-Blot® Turbo™, Bio-Rad, USA) at 25V for 30 minutes. Membranes were blocked with 5% non-fat milk in Tris-buffered saline containing 0.1% Tween-20 (TBST) for 1 hour at ambient temperature to minimize non-specific binding, and subsequently incubated with primary antibodies (**Supplementary Table 1**) overnight at 4°C with gentle rocking. Following three 10-minute TBST washes, membranes were incubated with appropriate horseradish peroxidase-conjugated secondary antibodies for 1 hour at ambient temperature. Immunoreactive bands were visualized by enhanced chemiluminescence using SuperSignal™ West Dura Extended Duration Substrate (Thermo Fisher Scientific, USA) and digitally captured on a ChemiDoc™ MP Imaging System (Bio-Rad, USA) with optimized exposure parameters. Densitometric analysis was performed using ImageJ software (NIH, version 1.53c) to quantify relative protein expression. β-actin (1:25,000; Sigma, USA) served as endogenous loading control for normalization of target protein signals. Western blot data are presented as fold change relative to sham controls.

### 2.9 Quantitative spatial analysis of biomarker distribution via immunofluorescent histochemistry

Tissue sections mounted on Polysine adhesion slides underwent immunofluorescent labeling according to established protocols.^40^ Following rehydration, sections were subjected to three 5-minute washes with tris-buffered saline containing 0.1% Tween-20 (TBS-T, pH 7.6, Sigma, USA) and subsequently blocked with 10% normal goat serum (Abcam, USA) supplemented with 1% bovine serum albumin (Fraction V, Sigma, USA) for 1 hour at ambient temperature to minimize non-specific binding. Primary antibody incubation (**Supplementary Table 2**) was performed in humidified chambers at 4°C for 16-18 hours. Following three TBS-T rinses, sections were incubated with appropriate fluorophore-conjugated secondary antibodies (1:400 dilution) for 1 hour at ambient temperature with light protection. Nuclear counterstaining was performed using 4’,6-diamidino-2-phenylindole (DAPI, 1:2000, Thermo Fisher Scientific, USA) for 10 minutes, followed by three additional TBS-T washes. Sections were mounted with Vectashield Antifade mounting medium (Vector Laboratories, USA) and coverslipped. Fluorescent micrographs were acquired using an EVOS M7000 imaging system (Thermo Fisher Scientific, USA) equipped with appropriate filter at 20× magnification. Exposure parameters were standardized within experimental series to ensure valid comparative analyses. Quantitative image analysis was performed using ImageJ software (version 1.53c, National Institutes of Health, Bethesda, USA) with consistent thresholding parameters applied across all specimens.

Microvessel density was assessed by calculating Lectin-positive area as a percentage of total analyzed area (0.25 mm²) within anatomically defined regions including CA1 pyramidal layer and hippocampal fissure. Immunoreactivity for endothelin-1, Iba1, GFAP, APP, and Aβ42 was quantified semi-automatically using standardized region-of-interest selection and background subtraction algorithms.^41^ Signal intensities were quantified with pixel values ranging from 0 (minimum) to 255 (maximum). Raw immunofluorescence data for microglial (Iba1), astrocytic (GFAP), and vasoactive (endothelin-1) markers are presented as reciprocal intensity (arbitrary units, AU). Endothelin-1/Lectin signal ratio was calculated as an indicator of vasoconstrictive propensity within the microvasculature. Microvascular morphology was assessed by measuring vessel diameter from Lectin-positive structures, while endothelin-1 expression was quantified exclusively on vascular elements. Differential analysis of APP and Aβ42 localization (vascular versus parenchymal) was accomplished via co-localization analysis with Lectin immunoreactivity, with vascular characterized according to outer vessel diameter. Neuronal viability was evaluated using NeuN immunoreactivity, with morphological assessment conducted by two independent observers blinded to experimental conditions. Results are expressed as percentage of neurons exhibiting aberrant morphology (nuclear pyknosis, cytoplasmic shrinkage, or dendrite fragmentation) relative to total neuronal count per field.

### 2.10 Quantification of microvascular constriction and collapse

To evaluate overall microvascular constriction and collapse, brain sections (18 μm thickness) were subjected to standard hematoxylin and eosin (H&E) staining protocol. Stained sections were visualized using an EVOS M7000 imaging system (Thermo Fisher Scientific, USA) at 20× magnification with consistent illumination and exposure parameters. Digital micrographs of anatomically defined regions were systematically acquired, focusing on the CA1 stratum pyramidale and hippocampal fissure. Parenchymal microvessels (<100 μm diameter) were identified based on characteristic morphological features including endothelial lining, smooth muscle layer, and identifiable luminal space. Total vessel counts were manually enumerated throughout each anatomical region by two independent observers blinded to experimental conditions, with agreement by consensus for discrepant classifications. A minimum of 100 vessels per region were analyzed across multiple tissue sections to ensure adequate sampling. Vascular collapse was rigorously determined using dual complementary criteria and expressed as percentage of total arteriole number within each region of interest. The primary criterion for identifying vascular collapse was the presence of perivascular space between internal and external elastic laminae, visualized as distinctive separation between the vascular wall layers. The secondary criterion was minimum 30% reduction in luminal diameter compared to morphologically normal vessels within the same anatomical region. This dual-criteria approach was implemented to minimize potential misclassification, as tissue processing-induced deformation artifacts could potentially manifest as artificial separation between vascular laminae, thus confounding accurate assessment when relying solely on perivascular space as a collapse indicator.^42^ Vessels meeting both criteria were classified as collapsed, while those meeting only one criterion underwent additional morphological evaluation before final classification. The percentage of collapsed microvessels was calculated using the formula: (number of collapsed vessels/total vessel count) × 100.

### 2.11 Statistical Analysis

Sample size determination was conducted a priori using statistical power analysis. Specifically, power calculations were performed using a two-sided 95% confidence interval for a single mean with a power (1-β) of 0.80, based on effect sizes observed in previous investigations employing comparable methodologies. This analysis determined that n=6 animals per experimental group was sufficient to detect statistically significant differences in Western blot analyses and immunofluorescent examinations. For behavioral assessments, n=9 animals per group was calculated as necessary to detect significant differences in novel object recognition and Y-maze task performance.

All statistical analyses were conducted using GraphPad Prism software (version 9, GraphPad Software Inc., USA). Data are presented as mean ± standard deviation (SD) unless otherwise specified, with statistical significance established at p<0.05. Prior to parametric testing, the normality of data distribution was verified using both Shapiro-Wilk and Kolmogorov-Smirnov tests. For datasets meeting normality assumptions, between-group differences in cognitive performance, neuropeptide expression, and vascular/non-vascular parameters were assessed using one-way analysis of variance (ANOVA) followed by Tukey’s post-hoc test to correct for multiple comparisons.

For longitudinal behavioral and biomarker assessments, repeated measures ANOVA was employed to evaluate time-dependent changes across experimental groups. In instances where normality assumptions were violated, non-parametric equivalents (Kruskal-Wallis test followed by Dunn’s multiple comparisons) were utilized. Correlation analyses were performed between behavioral metrics (novel object recognition and Y-maze performance) and molecular parameters (neuropeptide expression levels, vascular integrity markers, and non-vascular indicators) using Pearson’s correlation coefficient for normally distributed data or Spearman’s rank correlation for non-parametric datasets. These correlations were visualized as either scatter plots with linear regression or composite heat maps to illustrate relationship patterns across multiple variables simultaneously.

## 3. Results

### 3.1 Global proteomic profiling reveals distinct expression patterns in the hippocampus following CCH

To determine the potential pathways involved in CCH pathophysiology, samples from late-stage CCH (6 weeks post-2VO) were analyzed by proteomic methods. Principal component analysis (PCA) was conducted to evaluate sample relationships and identify primary sources of variation in the proteomic dataset. PCA revealed clear segregation between chronic CCH and sham groups (n=5) (**Fig. 1A**). Differential expression analysis identified 688 significantly altered proteins from a total of 2388 proteins. Of these, 546 proteins were upregulated (≥2 fold-change) and 142 were downregulated (≤0.3-fold change) (**Fig. 1B**). The volcano plot illustrates the distribution of protein expression differences between groups, with the logarithm base 2 of fold change on the horizontal axis and the negative logarithmic-transformed p-value on the vertical axis. Functional categorization of differentially expressed proteins revealed seven major protein classes: synaptic signaling (14%), vasomotricity function (34%), amyloid plaque formation (10%), coagulation cascade (4%), neurotransmission (8%), oxidative stress (20%), and inflammation signaling pathway (10%) (**Fig. 1C**). Protein-protein interaction (PPI) network analysis using STRING revealed extensive interconnectivity among proteins mediating vasomotor function, oxidative stress response, and amyloid-associated signaling cascades. Notably, synaptic signaling components demonstrated significant interactions exclusively with the vasomotor regulatory network, with neuronal nitric oxide synthase (NOS1) serving as a central hub in this interaction (**Fig. 1D**). Conversely, proteins involved in coagulation processes, inflammatory signaling, and neurotransmission displayed minimal interaction connectivity following CCH. Heat map visualization of differentially expressed proteins categorized by functional domains revealed significant alterations in essential regulatory molecules, particularly those mediating vasomotor responses. The substantial enrichment of vasomotor-associated proteins, coupled with their remarkably consistent directional changes in CCH condition (**Fig. 1E**), indicates a coordinated dysregulation of vascular tone modulatory pathways under pathological conditions. Collectively, these results indicate that vasomotricity-related proteins exhibited the most pronounced alterations, suggesting significant modifications in critical protein interactions and signaling pathways during CCH progression.

**Figure 1.**
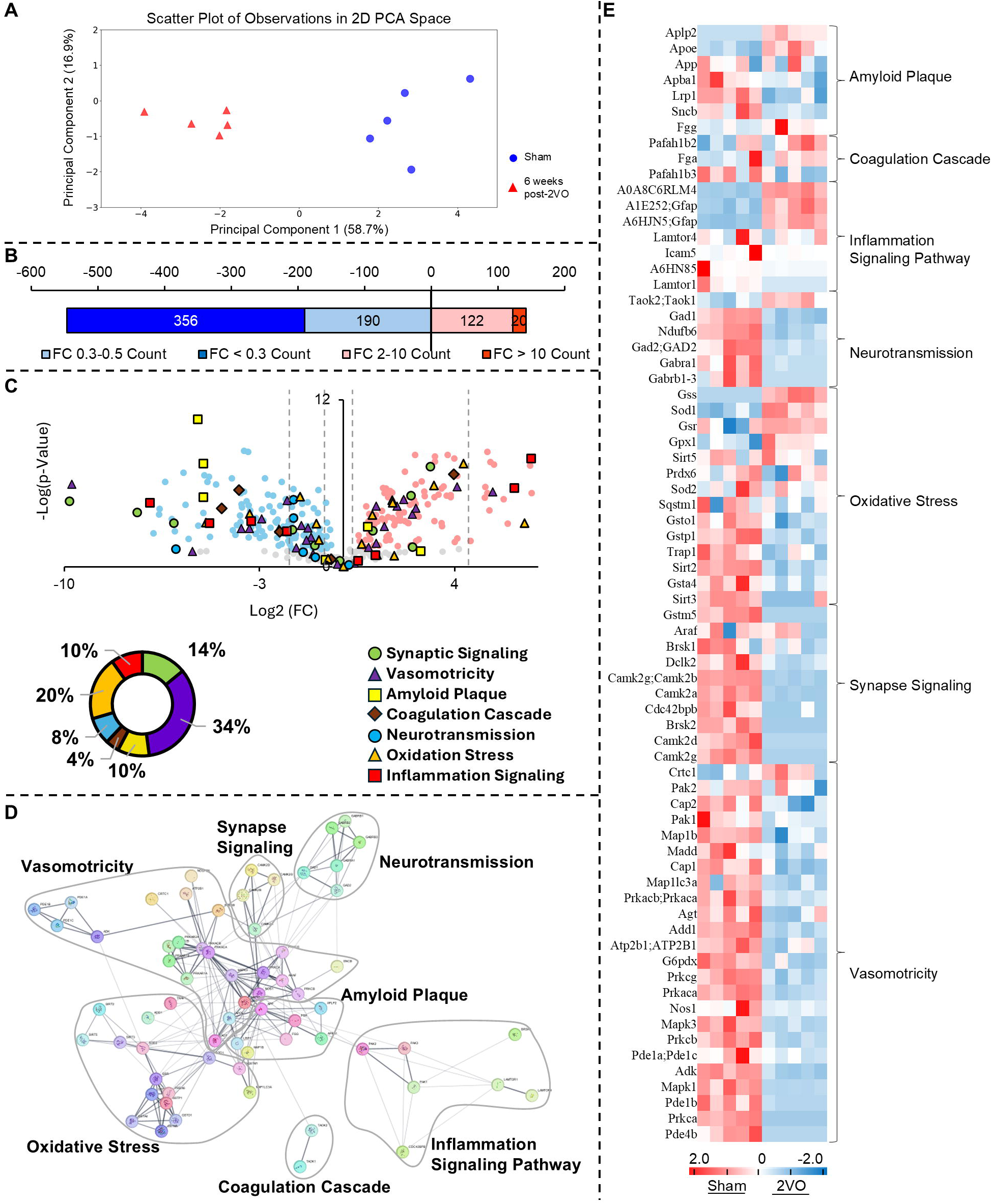
Comprehensive proteomic analysis reveals differential molecular signatures in hippocampal tissue following chronic cerebral hypoperfusion. **(A)** Principal Component Analysis revealing statistically significant separation (P<.05) between sham (blue) and CCH (red) experimental cohorts. **(B**) Quantitative distribution analysis of significantly modulated proteins (fold change thresholds: ≤0.3 and ≥2). **(C)** Volcano plot representation of differentially expressed proteins, comprising 546 significantly upregulated and 142 significantly downregulated proteins (fold change ≥ 2 or ≤ 0.3; P<.05). Categorical distribution diagram illustrating the percentage of differentially expressed proteins across functional domains: synaptic signaling, vasomotricity, amyloid plaque formation, coagulation cascade, neurotransmission, oxidative stress, and inflammatory signaling. **(D)** Protein-protein interaction network analysis of 71 differentially expressed proteins (nodes) organized into functional clusters. **(E)** Hierarchical clustering heatmap demonstrating differential expression patterns of 71 proteins between experimental groups (P<.05). (CCH: chronic cerebral hypoperfusion; PCA: principal component analysis; 2VO: bilateral common carotid artery occlusion)

### 3.2 CCH induces progressive microvascular deterioration in discrete hippocampal subfields

Vasoconstrictive responses and consequent morphological remodeling of the cerebral microvasculature were quantitatively assessed across three distinct stages of CCH: early (2 weeks post-2VO), middle (4 weeks post-2VO), and late (6 weeks post-2VO). Microvascular dynamics in the CA1 region and hippocampal fissure were evaluated through concurrent analysis of endothelin-1 immunoreactivity and Lectin-based vascular labeling (**Fig. 2A, Supplementary Fig. 2A**). Quantitative assessment revealed progressive intensification of perivascular endothelin-1 expression following CCH, indicating escalating vasoconstrictive signaling (**Fig. 2B-2E**). This increase was initially detectable at 2 weeks post-2VO and continued to amplify throughout the 6-week experimental timeline in both hippocampal regions. Notably, endothelin-1 upregulation demonstrated pronounced vessel-caliber specificity. Microvasculature within the CA1 region and smaller vessels (<20 μm diameter) within the hippocampal fissure exhibited the most substantial endothelin-1 expression at 2 weeks post-2VO, with continued progressive increases thereafter (**Fig. 2B, 2C**). This vessel-size dependent pattern suggests differential vulnerability of microvasculature to vasoconstrictive mechanisms during CCH progression.

**Figure 2.**
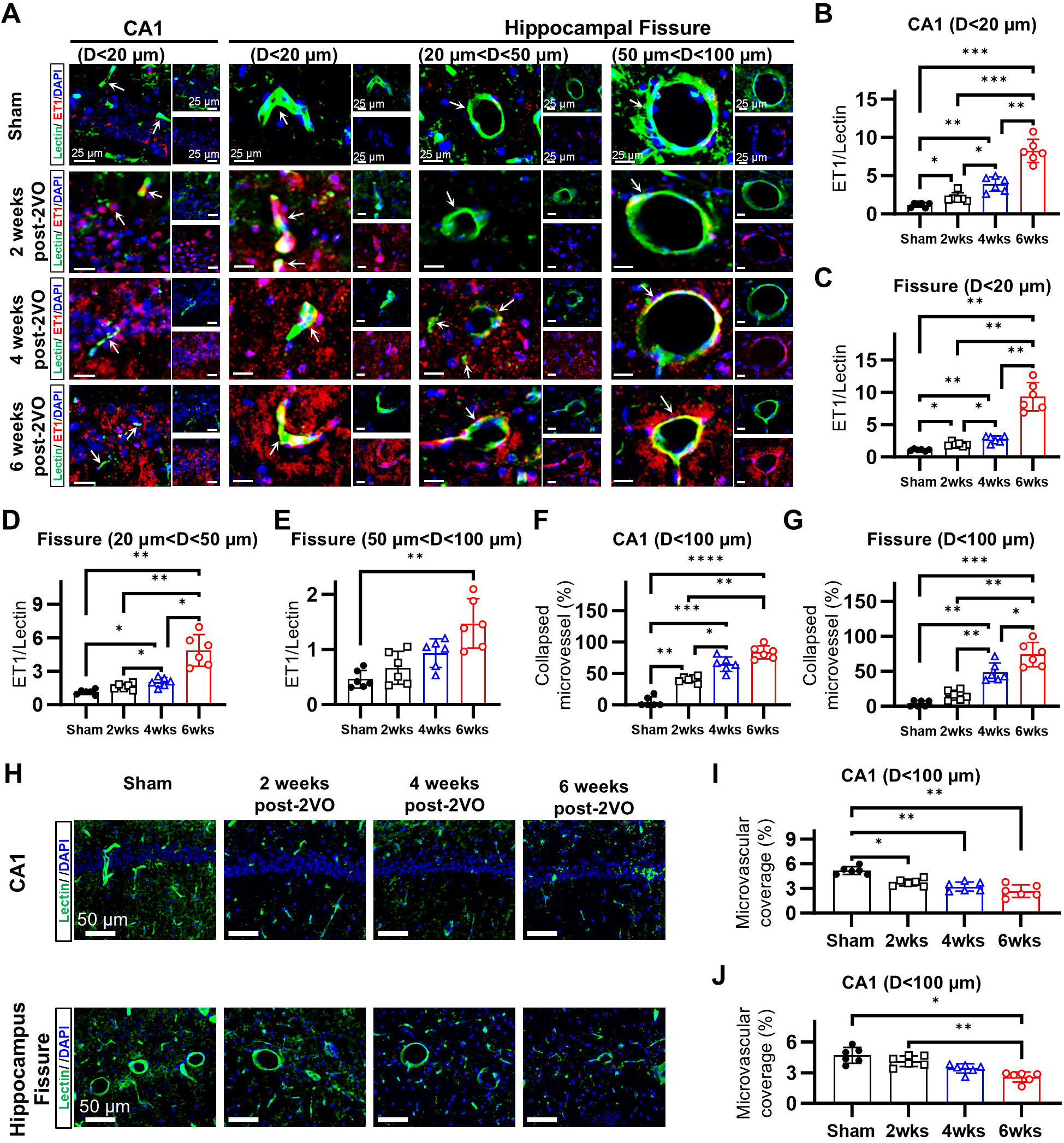
Chronic cerebral hypoperfusion induces progressive microvascular deterioration in hippocampal subfields. Temporal assessment of microvascular constriction and degeneration in the CA1 and hippocampal fissure of CCH-induced rats at 2, 4, and 6 weeks post-2VO via immunofluorescence analysis. **(A)** Immunofluorescent visualization of ET-1/Lectin co-localization within the CA1 region and hippocampal fissure demonstrating progressive microvascular constriction. **(B)-(E)** Quantitative analysis of ET-1/Lectin signal ratio showing continuous temporal elevation throughout the 6-week experimental timeframe, with vessel size-dependent vulnerability. **(F)-(G)** Morphometric analysis of vascular collapse demonstrating significant stepwise progression at 2, 4, and 6 weeks post-2VO compared to sham controls within both CA1 and hippocampal fissure regions. **(H)** Immunofluorescent visualization of ET-1 within the CA1 region and hippocampal fissure demonstrating progressive microvascular degeneration. **(I)-(J)** Quantitative assessment of Lectin-positive signal within the CA1 and hippocampal fissure showing progressive diminution, indicative of ongoing vascular degeneration. (CCH: chronic cerebral hypoperfusion; 2VO: bilateral common carotid artery occlusion; ET-1: endothelin-1; DAPI: 4’,6-diamidino-2-phenylindole; D: diameter; *p < 0.05, **p < 0.01, ***p < 0.001, ****p < 0.0001)

The functional consequences of enhanced endothelin-1 signaling were evident in the progressive microvascular constriction/collapse observed within the hippocampus (**Supplementary Fig. 2B**). Morphometric analysis revealed significant increases in the proportion of constricted vessels beginning at 2 weeks post-2VO, culminating in 83.5% of vessels exhibiting constriction in the CA1 region by 6 weeks (**Fig. 2F**). Similar patterns of progressive vasoconstriction were observed within the hippocampal fissure (**Fig. 2G**). Concurrent with these vasoconstrictive changes, we documented progressive diminution of lectin-positive vascular signals, indicating substantial microvascular degeneration (**Fig. 2H**). This rarefaction resulted in 59.2% reduction of vascular coverage in the CA1 region (**Fig. 2I**) and 50% reduction in the hippocampal fissure by 6 weeks post-2VO (**Fig. 2J**).

These findings demonstrate that CCH induces temporally progressive, vessel-caliber specific microvascular constriction, with the smallest vessels exhibiting the earliest and most pronounced vasoconstrictive alterations across the early, middle, and late stages of cerebral hypoperfusion. Furthermore, these vasoconstrictive changes precede and potentially contribute to subsequent microvascular degeneration. The vessel-size specificity and temporal progression of these alterations suggest that microvascular constriction represents a primary pathophysiological mechanism rather than a secondary consequence of CCH, potentially serving as a key initiating factor in the cascade of pathological events leading to hippocampal dysfunction.

### 3.3 Microvascular constriction demonstrates strong temporal correlation with progressive cognitive deterioration in CCH

To characterize the cognitive sequelae of CCH, we evaluated both short-term working recognition memory and long-term spatial memory utilizing novel object recognition and Y-maze paradigms, respectively. Our findings demonstrated that CCH induces progressive deterioration of short-term working recognition memory, as evidenced by steadily declining discrimination indices across the experimental timeline (sham: 0.34±0.22, 2 weeks post-2VO: −0.019±0.25, 4 weeks post-2VO: −0.14±0.23, 6 weeks post-2VO: −0.36±0.28, p<0.05) (**Fig. 3A**).

**Figure 3.**
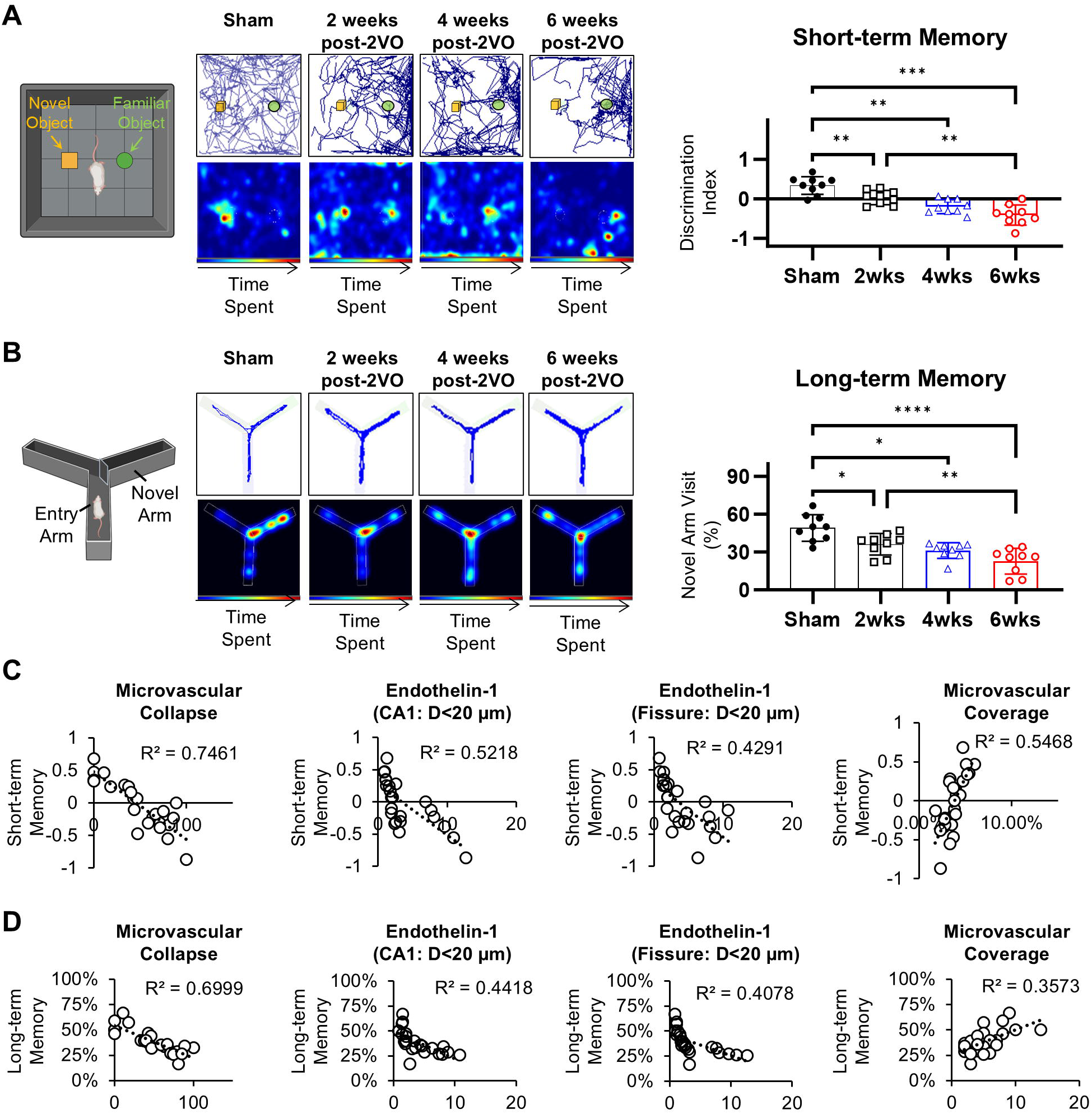
CCH-Induced Cognitive Impairment Demonstrates Significant Correlation with Microvascular Pathology, Particularly Microvascular Collapse. **(A)** NOR assessment demonstrating temporal progression of short-term working reference memory deficits at 2, 4, and 6 weeks post-2VO. **(B)** Y-maze spatial navigation paradigm revealing progressive deterioration of long-term spatial memory function across the experimental timeline (2, 4, and 6 weeks post-2VO intervention). **(C)** Regression analysis demonstrating relationship between microvascular pathological parameters (collapse and vascular degeneration) and short-term memory deficits. Short-term memory dysfunction exhibited significant correlation with microvascular collapse, with secondary correlation to vascular degeneration in hippocampal formations. **(D)** Regression analysis between microvascular pathological parameters (collapse and vascular degeneration) and long-term memory deficits. Long-term memory dysfunction displayed significant correlation with microvascular collapse, with secondary correlation to vascular degeneration in hippocampal formations. (CCH: chronic cerebral hypoperfusion; 2VO: bilateral common carotid artery occlusion; NOR: novel object recognition test; *p < 0.05, **p < 0.01, ***p < 0.001, ****p < 0.0001)

This progressive impairment was reflected by diminishing preferential interaction with novel objects relative to familiar objects. Concurrently, we observed analogous deterioration in long-term spatial memory function, demonstrated by progressive reduction in novel arm preference during Y-maze assessment (sham: 46.4%±14%, 2 weeks post-2VO: 36.9%±8%, 4 weeks post-2VO: 34.5%±9%, 6 weeks post-2VO: 27.2%±11%, p<0.05) (**Fig. 3B**). These decrements were characterized by diminishing preferential exploration of novel arms within the maze apparatus.

To determine the mechanistic relationship between cerebrovascular pathology and cognitive dysfunction, we conducted comprehensive correlation analyses examining correlations between various parameters of microvascular injury and cognitive performance metrics. Notably, microvascular constriction/collapse in the CA1 region demonstrated the strongest correlation with short-term working recognition memory deficits (R²=0.7461), establishing a robust relationship between structural microvascular integrity and cognitive function (**Fig. 3C**). Moderate correlations were observed between short-term memory impairment and endothelin-1 expression (CA1: R²=0.5218, hippocampal fissure: R²=0.4291), a marker of vasoconstriction, as well as with indices of vascular degeneration (R²=0.5468) (**Fig. 3C, Supplementary Fig. 3**). Similar associations were identified between microvascular parameters and long-term spatial memory performance, although correlation coefficients were generally lower than those observed for short-term memory outcomes (**Fig. 3D, Supplementary Fig. 3**). This differential correlation suggests potential domain-specific vulnerability to microvascular dysfunction, with short-term recognition memory processes demonstrating greater sensitivity to cerebrovascular compromise.

These findings collectively demonstrate that CCH induces progressive cognitive deterioration, with particularly pronounced effects on short-term working memory function. The strong correlations between cognitive performance metrics and parameters of microvascular dysfunction—especially microvascular collapse and endothelin-1-mediated vasoconstriction—provide compelling evidence for the primacy of cerebrovascular pathology in CCH-induced cognitive impairment. These data establish microvascular constriction/collapse as a critical determinant in both the initial manifestation and subsequent progression of cognitive decline in the context of CCH.

### 3.4 Vascular amyloidogenic protein expression, but not non-vascular or total levels, correlates with cognitive decline in CCH

To establish the primacy of microvascular pathology in cognitive deterioration during CCH, we analyzed temporal changes in amyloid-associated protein markers previously implicated in VCI and AD.^13^ Hippocampal tissue analysis revealed that CCH induces progressive increases in phosphorylated tau (pTau) and α-synuclein throughout the experimental timeline (**Fig. 4A-4B**). In contrast, amyloid precursor protein (APP) demonstrated non-linear kinetics, with expression increasing until 4 weeks post-2VO, followed by subsequent reduction at 6 weeks (**Fig. 4A-4B**). Correlation analyses examining the relationship between identified protein markers and cognitive performance metrics revealed modest associations for both memory domains assessed. Short-term memory performance demonstrated determination coefficients (R²) ranging from 0.1791 to 0.3992 (**Fig. 4C**), while long-term memory assessments yielded R² values between 0.2297 and 0.4587 (**Supplementary Fig. 4A-4C**). The limited explanatory power of these correlations suggests that whole-tissue abundance of these protein biomarkers may not constitute the principal mechanistic driver of cognitive dysfunction in this model system.

**Figure 4.**
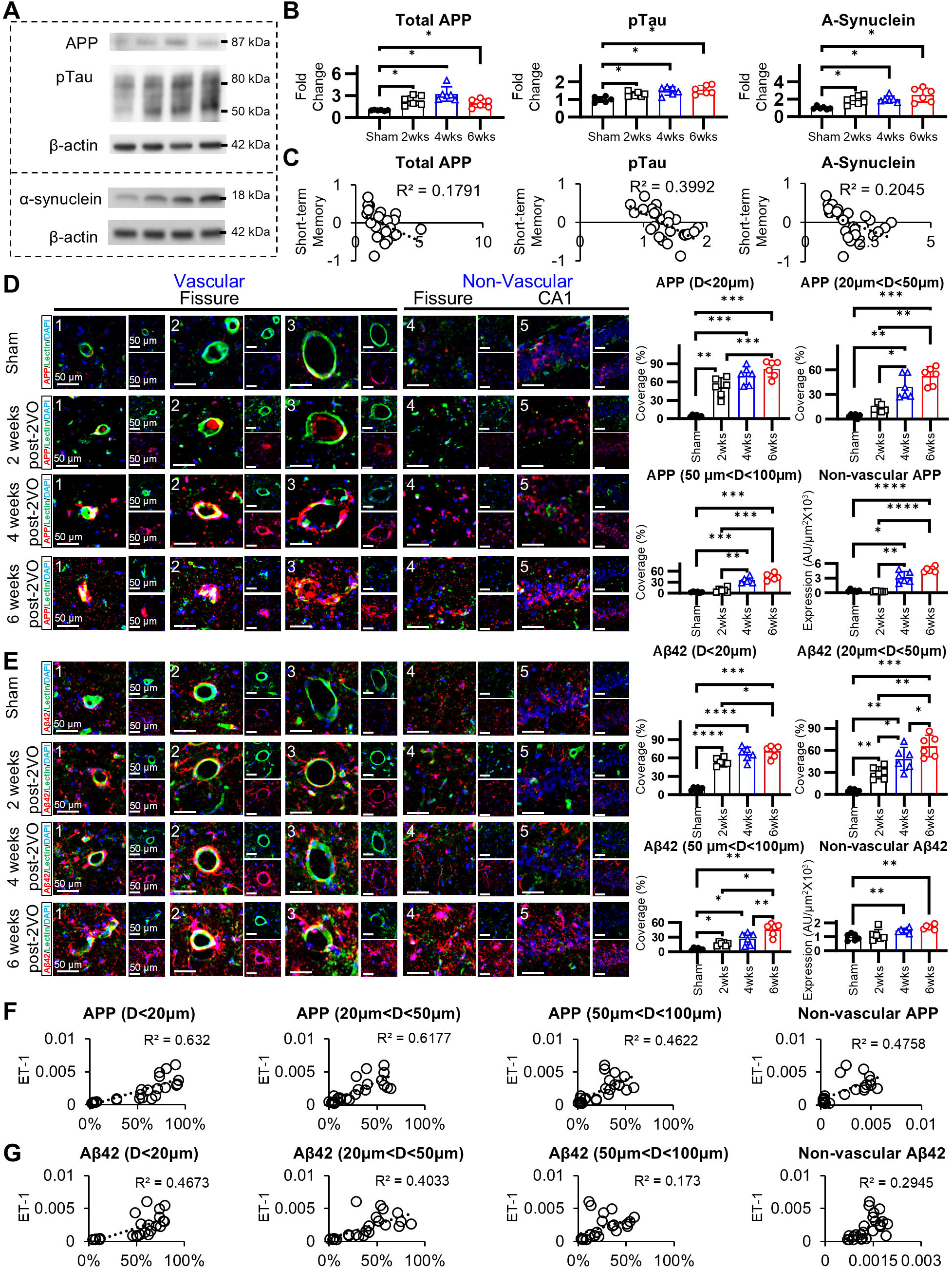
Vascular-associated, but not total, amyloidogenic protein expression correlates with cognitive impairment in chronic cerebral hypoperfusion. Temporal assessment of amyloidogenic proteins (APP, pTau, and α-synuclein) in CCH-induced rats at 2, 4, and 6 weeks post-2VO utilizing Western blot analysis and immunofluorescence techniques. **(A)-(B)** Densitometric analysis demonstrating progressive upregulation of APP (maximal at 4 weeks), pTau, and α-synuclein protein expression compared to sham controls. **(C)** Regression analyses between total amyloidogenic protein expression and short-term memory performance following CCH. Total APP, pTau, and α-synuclein expression demonstrate poor correlation with short-term memory deficits in CCH. **(D)-(E)** Differential quantification of vascular-associated and parenchymal APP and Aβ42 accumulation within the CA1 region and hippocampal fissure. Vascular-associated APP and Aβ42 deposition exhibits vessel size-dependent progression, with primary involvement of microvessels followed by larger caliber vessels. Significant parenchymal APP or Aβ42 accumulation manifests only after 4 weeks post-intervention, subsequent to vascular deposition. **(F)-(G)** Correlation analyses between APP/Aβ42 expression and ET1 immunoreactivity. Correlation coefficients demonstrate inverse relationship with vessel caliber, with highest correlation in microvessels and lowest in non-vascular regions. (CCH: chronic cerebral hypoperfusion; APP: amyloid precursor protein; pTau: phosphorylated tau; Aβ42: amyloid β42; ET-1: endothelin-1; *p < 0.05, **p < 0.01, ***p < 0.001, ****p < 0.0001)

Given these findings, we conducted detailed spatial analysis of amyloid distribution to determine whether vascular-specific amyloid deposition might demonstrate stronger associations with cognitive outcomes. Notably, immunohistochemical analysis revealed predominant accumulation of both APP and Aβ42 within vascular structures rather than non-vascular hippocampal regions ((**Fig. 4D-4E**, **Supplementary Fig. 4D**). Vessel size-specific analysis demonstrated distinct temporal patterns of amyloid accumulation, with microvasculature (<20 μm diameter) exhibiting the earliest increases in both APP and Aβ42, followed by progressive involvement of medium (20-50 μm) and large (>50 μm) vessels (**Fig. 4D-4E**, **Supplementary Fig. 4D**). Temporospatial analysis revealed that in larger vessels, APP was initially detectable within the luminal compartment at 2 weeks post-2VO, followed by perivascular aggregation at 4 weeks, and ultimately accumulation within surrounding parenchymal tissue by 6 weeks. Similar progression was observed with Aβ42, albeit with less pronounced immunoreactivity compared to APP. Importantly, non-vascular parenchymal and CA1 regional amyloid deposition demonstrated delayed kinetics relative to vascular accumulation, becoming detectable only after substantial vascular amyloid burden had developed.

To quantitatively assess the relationship between vascular amyloid deposition and microvascular dysfunction, correlation analyses were performed between amyloid markers and vessel constriction. Vascular APP accumulation strongly correlated with microvascular constriction-associated marker endothelin-1, particularly in the smallest microvessels (R²=0.632), with progressively lower correlations in medium microvessels (R²=0.618), large microvessels (R²=0.462), and non-vascular regions (R²=0.476) (**Fig. 4F**). Similarly, vascular Aβ42 demonstrated the strongest correlation with microvascular constriction in small vessels (R²=0.467), followed by medium vessels (R²=0.403), large vessels (R²=0.173), and non-vascular regions (R²=0.295) (**Fig. 4G**).

When examining correlations with cognitive outcomes, vascular amyloid deposition showed substantially stronger correlations compared to total tissue amyloid expression. Vascular APP demonstrated robust correlations with cognitive performance (R²=0.4777-0.5857), as did vascular Aβ42 (R²=0.3215-0.6262). In contrast, non-vascular APP showed weaker associations (R²=0.4345—0.4831), while non-vascular Aβ42 maintained moderate correlations (R²=0.3447-0.4189) (**Supplementary Fig. 4E-4F**).

These findings establish that vascular-specific amyloid deposition, particularly within the smallest constricted vessels, demonstrates the strongest correlations with both microvascular dysfunction and cognitive impairment. This spatial relationship provides compelling evidence that microvascular constriction precedes and potentially drives amyloid pathology, confirming the hypothesis that microvascular dysfunction plays a primary and causal role in CCH pathogenesis and resultant cognitive deterioration.

### 3.5 Pathogenic significance of early-onset and persistent vasoactive neuropeptide dysregulation in CCH

To elucidate the molecular mechanisms underlying microvascular dysfunction in CCH, we conducted systematic quantification of vasoactive neuropeptides implicated in CCH pathophysiology (**Fig. 5A-5G**). These neuropeptides were categorized according to their predominant pathophysiological functions: vasomotor regulation (endothelin-1, CGRP), amyloid pathology (high molecular-weight kininogen), coagulation (vasopressin, somatostatin [SST], somatostatin receptor 5 [SSTr5], hepcidin), vascular degeneration (angiotensin II receptor type 1 [AGTIIr1], angiotensin-converting enzyme [ACE]), oxidative stress (brain natriuretic peptide [BNP]), BBB integrity (pituitary adenylate cyclase-activating polypeptide [PACAP], adrenomedullin), and inflammatory processes (vasoactive intestinal peptide [VIP]). Classification was established according to primary and secondary functions documented in neurodegenerative conditions.^13,14^

**Figure 5.**
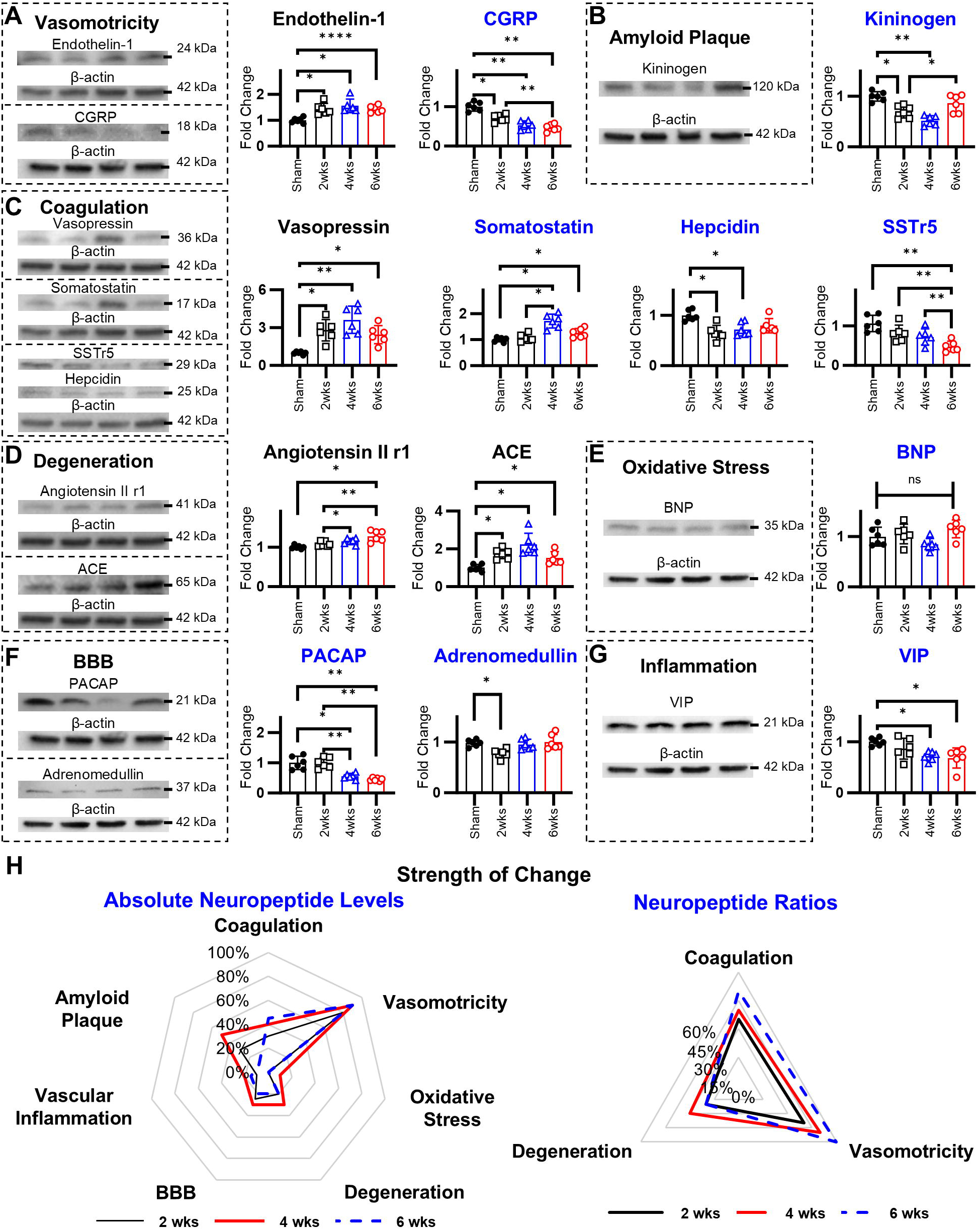
Chronic cerebral hypoperfusion induces progressive dysregulation of neuropeptides, predominantly those governing vasomotor function. Comprehensive temporal evaluation of neuropeptide expression in CCH-induced rats at 2, 4, and 6 weeks post-2VO using quantitative Western blot analysis. **(A)** Densitometric analysis demonstrating significant perturbation of vasomotor neuropeptides, characterized by upregulation of endothelin-1 and concurrent downregulation of CGRP. **(B)** Quantitative assessment revealing CCH-induced downregulation of kininogen, a neuropeptide susceptible to modulation by amyloid plaque formation. **(C)** Densitometric analysis of coagulation-regulatory neuropeptides demonstrating bidirectional dysregulation, with significant upregulation of vasopressin and somatostatin, contrasted by downregulation of hepcidin and SSTr5. **(D)** Quantitative assessment revealing CCH-mediated upregulation of neuropeptides implicated in vascular degeneration, including angiotensin II receptor 1 and ACE. **(E)** Densitometric analysis demonstrating significant downregulation of BNP, an oxidative stress-mitigating neuropeptide. **(F)** Quantitative assessment revealing CCH-induced downregulation of BBB-protective neuropeptides, including PACAP and adrenomedullin. **(G)** Densitometric analysis demonstrating significant downregulation of VIP, a neuropeptide integral to vascular inflammation regulation. **(H)** Comparative visualization depicting magnitude of alterations in absolute neuropeptide levels versus neuropeptide ratios. Absolute neuropeptide level analysis demonstrates predominant dysregulation of vasomotricity-regulating neuropeptides. Ratio analysis reveals CCH-induced shift in neuropeptide homeostasis toward a microenvironment deleterious to vasomotor function. (CCH: chronic cerebral hypoperfusion; SSTr5: somatostatin receptor 5; CGRP: calcitonin gene-related peptide; BNP: brain natriuretic peptide; ACE: angiotensin converting enzyme; PACAP: pituitary adenylate cyclase-activating polypeptide; VIP: vasoactive intestinal peptide; *p < 0.05, **p < 0.01, ***p < 0.001, ****p < 0.0001)

Temporal analysis revealed significant dysregulation across multiple neuropeptide systems following CCH. Vasomotor regulators demonstrated reciprocal alterations, with endothelin-1 exhibiting progressive upregulation while CGRP showed sustained downregulation throughout the 6-week observation period (**Fig. 5A**). High molecular-weight kininogen displayed biphasic regulation, with initial decreases at 2 and 4 weeks post-2VO followed by subsequent elevation at 6 weeks (**Fig. 5B**). Among coagulation-associated peptides, vasopressin—which functions both as a coagulation mediator and potent vasoconstrictor—demonstrated consistent upregulation throughout the experimental timeline. Somatostatin exhibited complex temporal dynamics with transient elevation at 4 weeks post-2VO, which was reversed at 6 weeks. Both hepcidin and SSTr5 showed sustained downregulation beginning at 2 weeks post-2VO (**Fig. 5C**). Vascular degeneration-associated neuropeptides displayed differential temporal patterns, with AGTIIr1 showing significant elevation exclusively at 6 weeks post-2VO, while ACE demonstrated progressive increases at 2 and 4 weeks followed by diminished expression at 6 weeks (**Fig. 5D**). BNP, an oxidative stress modulator, showed no significant alterations throughout the experimental course (**Fig. 5E**). Of the BBB and inflammation regulatory peptides, only PACAP (**Fig. 5F**) and VIP (**Fig. 5G**) exhibited significant decreases.

To evaluate the collective impact of these alterations, we calculated ratio analyses between protective and maladaptive neuropeptides across each pathomechanistic category. Notably, predominantly vasoconstrictive neuropeptides (endothelin-1 and vasopressin) demonstrated the most consistent and significant ratio decreases throughout the experimental timeline (**Supplementary Fig. 5A-5B**). While AGTIIr1 and ACE showed fewer significant ratio alterations, these were predominantly observed in relation to CGRP, PACAP, kininogen, and SSTr5, emphasizing the central importance of vasomotor regulatory peptides in CCH pathophysiology (**Supplementary Fig. 5A-5B**).

To determine the pathogenic significance of neuropeptide alterations in CCH, we conducted comprehensive correlation analyses examining both individual neuropeptides and neuropeptide ratios in relation to microvascular dysfunction and cognitive outcomes. Our findings revealed that neuropeptide dysregulation significantly impacts vasomotor function and coagulation parameters throughout CCH progression, as evidenced by both absolute expression levels and altered peptide ratios (**Fig. 5H**). Further correlation analyses between neuropeptide ratios and cognitive performance metrics demonstrated compelling associations with both short-term and long-term memory impairment (**Supplementary Fig. 5A-5B**). Particularly notable were ratios involving vasopressin, endothelin-1, SSTr5, CGRP, hepcidin, and VIP, which consistently demonstrated correlation coefficients (R²) exceeding 0.4 with memory performance measures. The strongest correlations were observed between CGRP:ET-1 ratio and novel object recognition performance (R²=0.62), followed by CGRP:vasopressin (R²=0.55), SSTr5:ET-1 (R²=0.44), and VIP:ET-1 (R²=0.51). While similar trends were observed with Y-maze performance, the correlation strengths were generally lower compared to those with NOR performance. When examining individual neuropeptides, ET-1, SSTr5, and CGRP demonstrated correlation coefficients with NOR exceeding 0.3, with CGRP exhibiting the strongest individual correlation (R²=0.83). This indicates a particularly prominent role for CGRP in maintaining cognitive function during cerebral hypoperfusion.

These findings demonstrate that CCH induces early and persistent dysregulation of vasoactive neuropeptides, which appears to be a primary pathophysiological mechanism underlying microvascular dysfunction and progressive cognitive deterioration. The strength and consistency of these correlations provide robust evidence supporting the concept that vasoactive neuropeptide imbalance serves as a principal driver of cognitive decline in CCH, rather than merely representing a secondary consequence of cerebrovascular insufficiency.

### 3.6 Non-vascular pathologies are not primary determinants of cognitive impairment in CCH

A comprehensive panel of markers associated with CCH pathophysiology was evaluated, encompassing BBB integrity, inflammatory processes, oxidative stress, coagulation cascade, angiogenic/anti-angiogenic signaling, vasomotor function, and endothelial cell dysfunction. BBB-associated proteins exhibited biphasic changes, with MMP-2 and occludin levels progressively increasing through 4 weeks post-2VO, followed by reversal at 6 weeks post-2VO (**Fig. 6A**). Inflammatory adhesion molecules displayed temporal divergence, with ICAM1 significantly elevated at 4 weeks post-2VO before declining at 6 weeks, while VCAM1 demonstrated significant upregulation exclusively at 6 weeks post-2VO (**Fig. 6B**). Oxidative stress markers, including SOD1 and nitrotyrosine, exhibited delayed responses, with significant elevations in nitrotyrosine observed starting at 4 weeks post-2VO and SOD1 increasing at the 6-week post-2VO timepoint (**Fig. 6C**). The coagulation factor fibrinogen showed early elevation at 2 and 4 weeks post-2VO, subsequently decreasing at 6 weeks post-occlusion (**Fig. 6D**). PDGFrβ expression demonstrated a transient increase at 2 weeks post-2VO, followed by progressive reduction, consistent with previously documented patterns of vascular degeneration (**Fig. 6E**). Endothelial function was markedly altered; eNOS expression exhibited progressive decline beginning at 2 weeks post-2VO (**Fig. 6F**). Vasomotor regulation was significantly altered, with marked reduction in the vasodilatory protein Pde1b, particularly at 4 weeks post-2VO (**Fig. 6G**) but no significant change of the vasoconstrictor regulator CRTC1 at 6 weeks post-2VO. Glial responses were characterized by non-significant increases in microglial activation, while astrocyte reactivity demonstrated progressive intensification from 2 through 6 weeks post-2VO, indicating sustained neuroinflammatory processes during CCH progression (**Fig. 6H, Supplementary Fig. 6A**).

**Figure 6.**
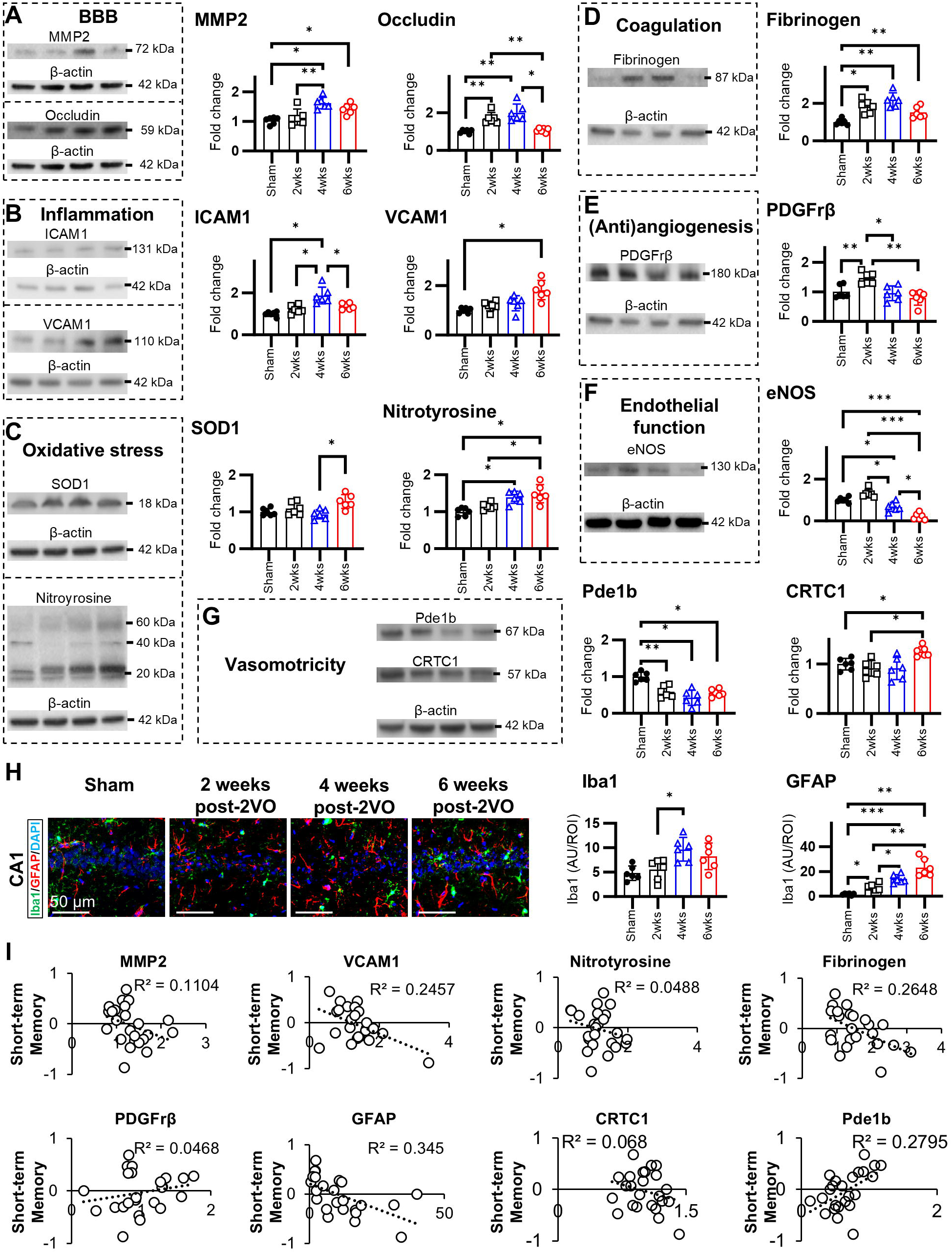
Non-vascular pathologies demonstrate limited contribution to cognitive dysfunction in chronic cerebral hypoperfusion. **(A)** Densitometric analysis revealing temporal dysregulation of MMP2, with significant elevation only at 4 weeks post-occlusion, while occludin expression increases at 2 and 4 weeks post-2VO. **(B)** Quantitative assessment demonstrating differential upregulation of adhesion molecules, with ICAM1 expression increasing exclusively at 4 weeks post-2VO, whereas VCAM1 expression elevates only at 6 weeks post-occlusion. **(C)** Molecular quantification revealing modest upregulation of oxidative stress markers SOD1 and NT at 6 weeks post-2VO. **(D)** Densitometric analysis demonstrating significant fibrinogen upregulation at 2 and 4 weeks post-2VO. **(E)** Quantitative assessment revealing biphasic regulation of PDGFrβ, characterized by transient elevation at 2 weeks post-2VO followed by significant reduction at 4 and 6 weeks post-2VO. **(F)** Molecular quantification demonstrating CCH-induced endothelial dysfunction markers. **(G)** Densitometric analysis revealing CCH-mediated downregulation of Pde1b concurrent with upregulation of CRTC1, representing opposing contributors to vasodilatory and vasoconstrictive signaling pathways, respectively. **(H)** Immunohistochemical quantification demonstrating predominant astrocytic activation within the CA1 region. **(I)** Correlation analysis between non-vascular pathological parameters and cognitive performance metrics revealing limited significant associations, with only four parameters demonstrating statistically significant correlation with cognitive outcomes. (CCH: chronic cerebral hypoperfusion; 2VO: bilateral common carotid artery occlusion; MMP2: matrix metalloproteinase 2; BBB: blood-brain barrier; ICAM1: intercellular adhesion molecule 1; VCAM1: vascular cell adhesion molecule 1; SOD: superoxide dismutase; eNOS: endothelial nitric oxide synthase; PDGFrβ: platelet derived growth factor receptor β; Pde1b: Phosphodiesterase 1B; CRTC1: CREB-regulated transcription coactivator 1; Iba1: ionized calcium binding adaptor molecule 1; GFAP: glial fibrillary acidic protein; *p < 0.05, **p < 0.01, ***p < 0.001, ****p < 0.0001)

Correlation analysis between non-vascular pathological markers and cognitive performance metrics revealed that only four parameters demonstrated statistically significant associations with cognitive outcomes. Among these, GFAP exhibited moderate correlations with both novel object recognition (R²=0.345) (**Fig. 6I**) and Y-maze performance (R²=0.3503) (**Supplementary Fig. 6C**), while Pde1b showed variable correlations with novel object recognition (R²=0.2795) (**Fig. 6I**) and Y-maze performance (R²=0.4675) (**Supplementary Fig. 5F**). VCAM1 and fibrinogen demonstrated weaker correlations compared to these two markers (**Fig. 6I, Supplementary Fig. 6C, 6E**). Temporal analysis of pathological progression revealed a biphasic pattern, with endothelial dysfunction and BBB disruption manifesting during the early phase CCH, while oxidative stress indices and astrocytic activation became prominent during later disease stages. Importantly, non-neuropeptide markers collectively demonstrated relatively weak correlations with both short-term and long-term memory parameters.

These findings indicate that non-neuropeptide, non-vascular pathological markers likely represent secondary consequences of CCH-induced pathophysiology rather than primary drivers of cognitive impairment. The limited correlative strength between these parameters and cognitive outcomes suggests that alternative mechanisms, particularly those involving vasoactive neuropeptides, may be more fundamentally involved in the pathogenesis of CCH-associated cognitive dysfunction.

### 3.7 Dysregulation of vasoactive neuropeptides as a primary mechanistic driver in CCH Pathophysiology

Comparative analysis of early-stage (2 weeks post-2VO) sensitivity markers, encompassing both neuropeptide and non-neuropeptide factors, was conducted to identify the principal pathogenic drivers in CCH (**Fig. 7A**). The results demonstrated that neuropeptides regulating vasomotricity function play a predominant role across the seven major pathomechanistic domains of CCH at this initial phase. Correlation analyses between neuropeptide and non-neuropeptide parameters revealed that dysregulation of vasoactive neuropeptides, particularly CGRP and endothelin-1, exhibited the strongest and most consistent associations with both short-term and long-term memory deficits. Inflammatory and oxidative stress biomarkers demonstrated moderate correlations specifically with long-term memory impairment (**Fig. 7B**).

**Figure 7.**
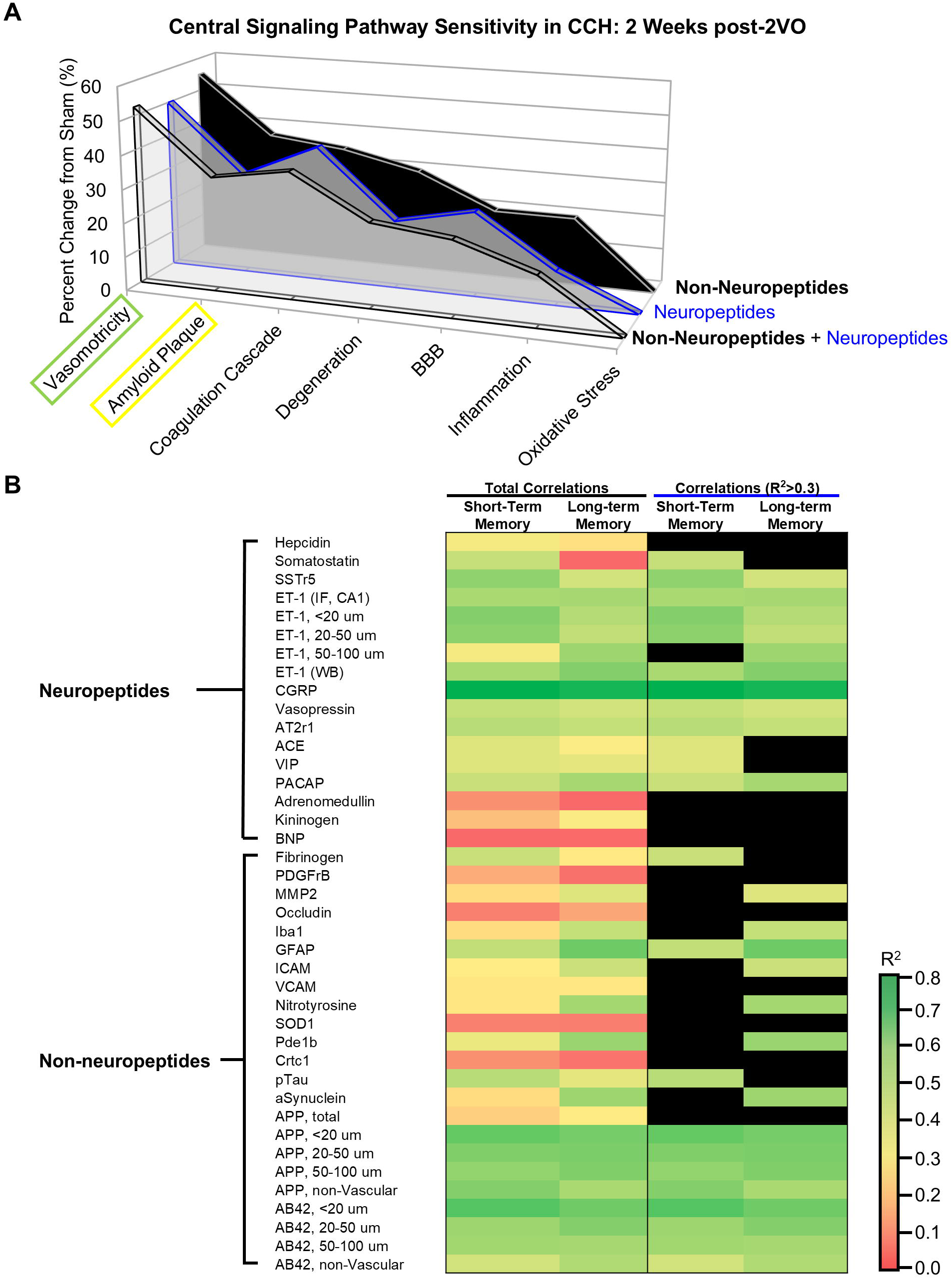
Vasoactive neuropeptides demonstrate superior sensitivity as biomarkers in chronic cerebral hypoperfusion. **(A)** Comparative analysis of neuropeptide versus non-neuropeptide markers across pathophysiological categories induced by CCH. Early vasomotor regulatory dysfunction exhibits predominant modulation by neuropeptidergic mechanisms rather than non-neuropeptide signaling pathways. **(B)** Hierarchical clustering heatmap depicting correlation coefficients between neuropeptide and non-neuropeptide markers with cognitive performance metrics following CCH induction. Neuropeptide-based markers demonstrate consistent and statistically significant correlation with both short-term and long-term memory impairment subsequent to CCH. Among the neuropeptides dysregulated by CCH, the vasodilatory neuropeptide CGRP exhibits the most robust correlation with both short-term and long-term memory deterioration (R² > 0.5). (CCH: chronic cerebral hypoperfusion; 2VO: bilateral common carotid artery occlusion; HEP: hepcidin; SST: somatostatin; SSTr5: somatostatin receptor 5; ET-1: endothelin-1; CGRP: calcitonin gene-related peptide; Vaso: vasopressin; AT2r1: angiotensin II receptor 1; ACE: angiotensin converting enzyme; VIP: vasoactive intestinal peptide; PACAP: pituitary adenylate cyclase-activating polypeptide; ADR: adrenomedullin; KIN: kininogen; PDGFrβ: platelet derived growth factor receptor β; MMP2: matrix metalloproteinase 2; Iba1: ionized calcium binding adaptor molecule 1; GFAP: glial fibrillary acidic protein; ICAM1: intercellular adhesion molecule 1; VCAM1: vascular cell adhesion molecule 1; SOD1: superoxide dismutase 1; pTau: phosphorylated tau; *p < 0.05, **p < 0.01, ***p < 0.001, ****p < 0.0001)

These findings establish that neuropeptides, especially those governing vascular tone and microcirculatory function, are critical mediators in CCH pathophysiology. The significant and consistent correlations between vasoactive neuropeptide alterations and cognitive deterioration across multiple domains suggest that these molecules function as fundamental pathogenic drivers in CCH progression. This evidence indicates that vasoactive neuropeptide dysregulation constitutes a primary mechanism underlying microvascular dysfunction and subsequent cognitive decline in vascular cognitive impairment, providing potential targets for therapeutic intervention.

### 3.8 CGRP supplementation preserves microvascular integrity and improves cognitive outcomes

CGRP functions as a principal vasoactive neuropeptide with significant implications in CCH pathogenesis. To investigate potential therapeutic interventions, we employed two distinct CGRP supplementation methodologies in CCH-induced animals: endogenous augmentation through DR and exogenous administration via intranasal delivery. Both supplementation strategies significantly enhanced cognitive performance, with approximately 80% improvement in short-term working recognition memory (**Fig. 8A, Supplementary Fig. 7A**) and significant enhancement in long-term spatial memory (**Fig. 8B, Supplementary Fig. 7B**). Molecular analyses revealed that both approaches elevated CGRP expression levels in CCH-affected hippocampus, with exogenous administration demonstrating superior efficacy (**Fig. 8C**). Additionally, SSTr5 expression was upregulated following both interventions, with endogenous CGRP supplementation exhibiting a more pronounced effect. Notably, CGRP supplementation attenuated endothelin-1 expression across treatment groups (**Fig. 8C**). Regarding pathological markers, both approaches significantly diminished the expression of APP and phosphorylated tau protein levels (**Supplementary Fig. 7C**). This concurrent reduction in critical neuropathological substrates suggests potential disease-modifying effects of CGRP-based interventions in the context of CCH.

**Figure 8.**
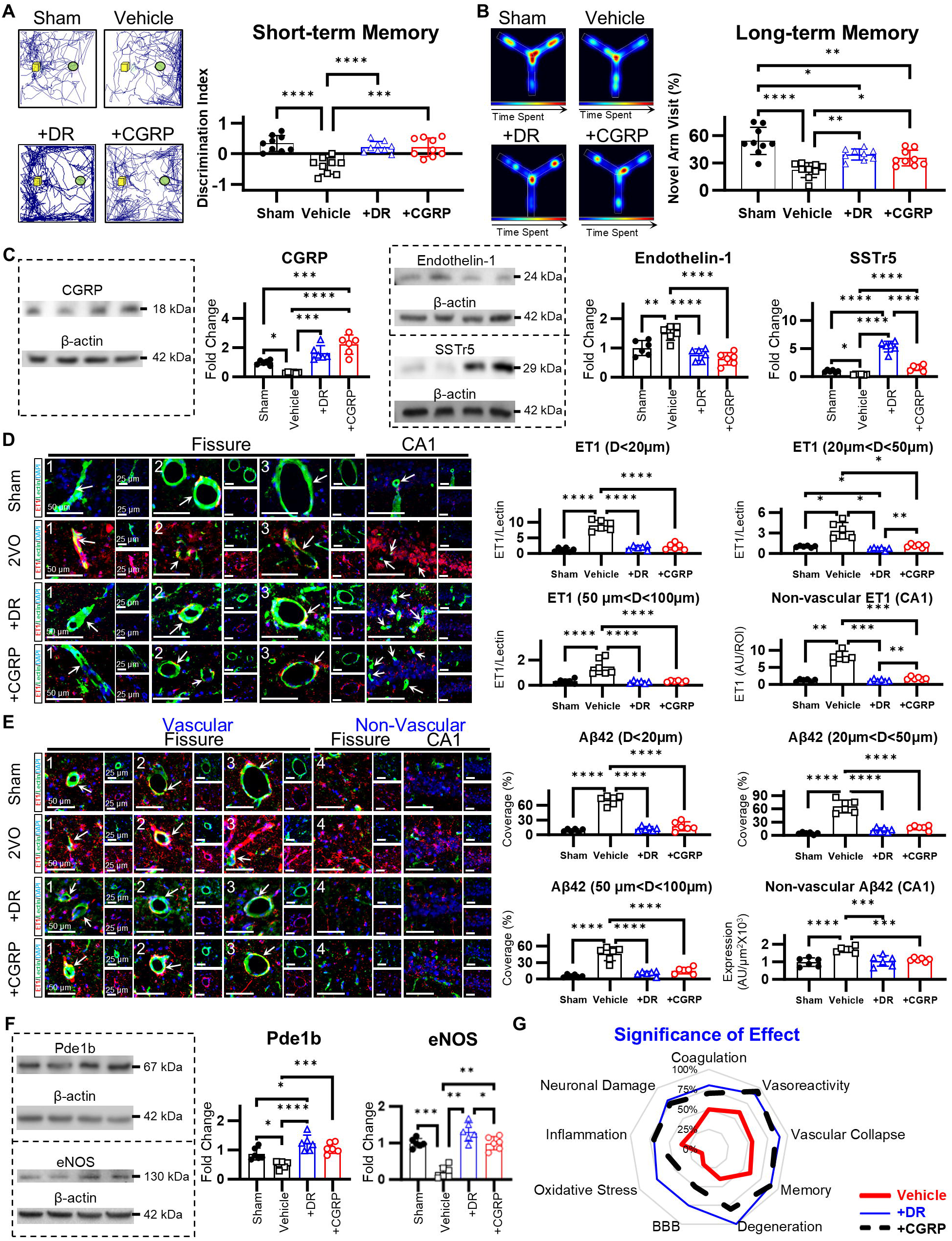
Vasoactive neuropeptide CGRP supplementation enhances cognitive function through attenuation of microvascular constriction and amyloidogenic pathology. **(A)-(B)** Quantitative assessment demonstrating endogenous and exogenous CGRP modulation ameliorates both short-term and long-term memory deficits at 6 weeks post-2VO. **(C)** Densitometric analysis revealing endogenous and exogenous CGRP modulation upregulates CGRP and SSTr5 expression while concurrently downregulating total ET-1 at 6 weeks post-2VO. **(D)** Immunofluorescence quantification demonstrating endogenous and exogenous CGRP supplementation attenuates ET-1 expression at 6 weeks post-2VO. **(E)** Quantitative immunohistochemical analysis showing endogenous and exogenous CGRP modulation reduces Aβ42 accumulation within the microvasculature of CA1 and hippocampal fissure regions at 6 weeks post-2VO. **(F)** Molecular quantification demonstrating CGRP supplementation significantly upregulates expression of Pde1b and eNOS. **(G)** Schematic representation illustrating differential therapeutic efficacy of endogenous versus exogenous neuropeptide modulation at 6 weeks post-2VO. Endogenous neuropeptide modulation via diving reflex activation demonstrates broader therapeutic impact across multiple CCH pathophysiological domains compared to exogenous CGRP administration. (CCH: chronic cerebral hypoperfusion; 2VO: bilateral common carotid artery occlusion; DR: diving reflex; CGRP: calcitonin gene-related peptide; SSTr5: somatostatin receptor 5; ET-1: endothelin-1; APP: amyloid precursor protein; pTau: phosphorylated tau; Pde1b: phosphodiesterase 1B; eNOS: endothelial nitric oxide synthase; *p < 0.05, **p < 0.01, ***p < 0.001, ****p < 0.0001)

Administration of both endogenous and exogenous CGRP significantly attenuated microvascular constriction, with endogenous delivery exhibiting superior vasodilatory efficacy (**Fig. 8D, Supplementary Fig. 7D**). The therapeutic effects of CGRP were observed across vessels of all calibers, including microvessels <20µm in diameter that demonstrated particular vulnerability to CCH. Additionally, CGRP intervention reduced parenchymal and pial arteriolar collapse (**Supplementary Fig. 7E**). Endogenous CGRP supplementation enhanced microvascular density within the CA1 hippocampal region to levels exceeding those observed in sham controls (**Supplementary Fig. 7F**). Analysis of amyloid pathology revealed that both CGRP delivery methods decreased vascular and parenchymal aggregation of Aβ42 (**Fig. 8E, Supplementary Fig. 7G**) in CCH. The reduction in vascular amyloid deposition was particularly pronounced in microvessels, suggesting that CGRP confers protection to the cerebral microvasculature most susceptible to hypoperfusion-induced damage. Comparative analysis of neuroinflammatory responses demonstrated that endogenous CGRP supplementation more effectively reduced astroglial activation relative to exogenous administration (**Supplementary Fig. 7H**). Moreover, endogenous and exogenous CGRP significantly mitigated inflammatory markers, oxidative stress indicators, and coagulation factors (p<0.05), and significantly increased pericyte (PDGFrβ) expression, affecting angiogenic processes (**Supplementary Fig. 7I**). Both supplementation strategies upregulated Pde1b and eNOS expression, with endogenous delivery eliciting more robust increases (**Fig. 7F**). Furthermore, both interventions significantly reduced neuronal degeneration in the CA1 region (**Supplementary Fig. 7J**). Comprehensive evaluation demonstrated that endogenous CGRP supplementation, compared to exogenous delivery, provided more extensive therapeutic benefits across multiple pathological domains, including microvascular dysfunction, amyloid accumulation, and non-vascular injury mechanisms following CCH (**Fig. 8H**).

The present findings demonstrate that CGRP administration effectively ameliorates microvascular constriction across all vessel calibers, with particularly pronounced effects in small-diameter vessels (<20µm) that exhibit heightened vulnerability in CCH. CGRP intervention significantly attenuated both vascular and parenchymal amyloid protein accumulation throughout hippocampal structures, indicating multifaceted effects on neuropathological progression. Furthermore, CGRP supplementation, especially via endogenous delivery mechanisms, significantly reduced non-vascular injury markers associated with CCH pathology. The intervention promoted endothelial cell recovery and maintained neuronal integrity during advanced stages of cerebral hypoperfusion. Collectively, these data indicate that CGRP plays a fundamental role in CCH pathophysiology, as its supplementation effectively counteracts multiple pathological processes, including microvascular dysfunction, endothelial cell deterioration, and amyloid protein aggregation, thereby attenuating consequent non-vascular tissue injury. These comprehensive neuroprotective effects establish CGRP as a potential therapeutic candidate addressing multiple pathological mechanisms in cerebral hypoperfusion-related neurodegeneration.

## 4. Discussion

This study provides the first evidence, to our knowledge, that dysregulation of vasoactive neuropeptides drives the pathogenesis of CCH and cognitive decline through microvascular constriction, initiating both vascular and non-vascular cascades characteristic of VCI. Among the investigated pathomechanisms, neuropeptides emerged as more significant contributors than non-neuropeptide factors, exhibiting stronger correlations with cognitive decline. Specifically, vasomotor-regulating neuropeptides, such as CGRP (associated with vasodilation) and endothelin-1 (associated with vasoconstriction), demonstrated the most robust correlations with cognitive impairment across various stages of CCH, identifying them as key mediators of CCH pathogenesis. Furthermore, the stronger correlations observed for CGRP compared to endothelin-1 suggests that impaired vasodilation has a greater impact on cognitive decline than enhanced vasoconstriction. Our data reveal a novel pathological cascade in CCH, wherein early vasodilatory impairment appears to initiate a pathological cascade encompassing microvascular coagulation, BBB disruption, neurodegeneration, and amyloid aggregation. Strikingly, among all non-neuropeptide markers evaluated, vascular amyloid aggregation demonstrated the strongest correlation with vasoconstriction and cognitive impairment, revealing a previously unrecognized link to CCH pathology. In contrast, classical non-vascular mediators, including oxidative stress and neuroinflammation, were observed exclusively during late CCH stages, suggesting their dependence on pre-existing microvascular dysfunction. CGRP supplementation significantly attenuated cognitive deterioration through prevention of microvascular constriction and subsequent reduction of both microvascular and non-vascular pathology. Notably, endogenous CGRP augmentation via DR demonstrated superior therapeutic efficacy compared to exogenous supplementation, potentially attributable to concurrent modulation of multiple vasoactive neuropeptides.

Our data indicate a predominant pathogenic role for neuropeptides compared to non-neuropeptide factors in CCH. Proteomic analysis revealed seven significantly dysregulated molecular pathways in late CCH, enabling comparative assessment of neuropeptide versus non-neuropeptide mediators within these identified networks. Systematic evaluation revealed that neuropeptide markers consistently exhibited stronger correlations with cognitive function metrics and demonstrated more pronounced, consistent alterations from baseline values relative to their non-neuropeptide counterparts within the same pathways. These findings align with established evidence demonstrating that ANS dysfunction correlates with cognitive decline severity^6,7^ and contributes to CCH and VCI through recurrent, transient orthostatic hypoperfusion episodes,^4–8^ potentially mediated through ANS-related neuropeptide dysregulation.^8,9,43,44^ Preclinical CCH investigations suggest neuropeptide dysregulation contributes to cognitive deterioration,^15–17^ and our findings further demonstrate that vasoactive neuropeptides modulate key pathomechanisms identified through proteomic analysis. Additionally, while the assessed non-neuropeptide markers have been previously proposed as potential VCI biomarkers,^45^ our data demonstrate substantially weaker correlations with cognitive outcomes compared to neuropeptide markers. This observation, combined with the established mechanistic role of neuropeptides in CCH pathophysiology, suggests that neuropeptides not only function as principal drivers of CCH pathogenesis relative to the investigated non-neuropeptide markers but may additionally serve as upstream regulators of their activity. This hierarchical relationship between neuropeptide dysregulation and downstream molecular alterations provides critical insights into the pathogenic cascade of CCH and identifies specific neuropeptide signaling pathways as potential high-priority therapeutic targets for intervention in VCI.

Endothelin-1 and CGRP, representing the most potent vasoconstricting and vasodilating neuropeptides respectively,^46,47^ emerge as critical mediators in CCH-associated microvascular dysfunction and cognitive deterioration. Clinical studies demonstrate direct involvement of these neuropeptides in VCID pathogenesis,^48–50^ with elevated endothelin-1 correlating with increased VCID risk and severity,^49,51^ while reduced CSF CGRP levels associate with AD.^50^ Mechanistically, both neuropeptides regulate CaMKII,^52,53^ a crucial vasoreactivity mediator previously identified through proteomic analysis as a key CCH pathology regulator.^54^ Our investigation revealed consistent endothelin-1 upregulation in hippocampal microvessels alongside progressive CGRP downregulation throughout CCH progression, corresponding with dysregulation of less potent neuropeptides, creating a proconstrictive microenvironment. Notably, CGRP demonstrated strongest correlations with microvascular dysfunction, suggesting impaired vasodilation may represent a more significant pathomechanism than enhanced vasoconstriction in CCH progression. As regulators of cerebral vascular tone^55^ implicated in intermittent orthostatic hypotension^8,43,44,56^ and hypoperfusion severity,^16^ these neuropeptides may function both as CCH pathogenesis drivers and predictive biomarkers—a distinctive characteristic compared to conventional VCI biomarkers that primarily predict severity rather than onset or progression.^45^

CCH manifests through multiple microvascular dysfunction mechanisms, notably BBB compromise and angiogenic dysregulation.^57^ Our findings, however, reveals that early, progressive microvascular constriction serves as the primary initiating event that triggers a cascade of subsequent microvascular pathologies. Vasoconstrictive neuropeptide dysregulation during initial CCH stages drives significant hippocampal vasoconstriction, preceding BBB compromise and vascular degeneration. This sequence has significant clinical implications, as neurovascular coupling impairment and capillary stalling strongly correlate with cognitive deficits in AD^58^ and cerebral small vessel disease—a recognized vascular dementia precursor.^59^ While previous investigations have implicated BBB dysfunction, angiogenesis, and coagulation in CCH pathogenesis,^25–28,60^ our temporal analysis demonstrating that microvascular constriction precedes these pathologies suggests its role as a predisposing factor for their development. Mechanistically, sustained vasoconstriction facilitates angiogenesis and BBB disruption through regional CBF reduction, which triggers pericyte recruitment to the affected region,^46,61,62^ as evidenced by our PDGFRβ expression observations. This pathophysiological sequence culminates in the formation of structurally compromised, permeable microvessels that are susceptible to collapse and subsequent degeneration—a process that propagates throughout the vascular network.^46,61,62^ The observed delayed MMP2 elevation coincides with increased BBB permeability,^62^ suggesting its occurrence subsequent to microvascular constriction. Notably, the progressive and sustained vasoconstriction demonstrated strongest correlation with cognitive decline compared to the plateaued/decreased developmental patterns and weaker correlations observed with MMP2 and fibrinogen. These findings collectively identify microvascular constriction as the factor most robustly associated with CCH pathogenesis and cognitive deterioration, highlighting its significance as a primary therapeutic target for intervention. The sequential nature of microvascular pathology development suggests that early therapeutic targeting of vasoconstriction may prevent the subsequent cascade of microvascular dysfunction that leads to irreversible vascular degeneration and cognitive impairment.

Previous VCI investigations have predominantly examined total/non-vascular amyloid^63^, while the spatiotemporal dynamics and pathogenic significance of vascular amyloid aggregation remain inadequately characterized. Our data demonstrate that vascular amyloid accumulation temporally precedes and directly facilitates non-vascular amyloid production and aggregation. This non-vascular amyloid deposition manifests exclusively during late/severe CCH stages, occurring only after vascular amyloid extends from the lamina to vascular walls, ultimately precipitating vascular collapse. Notably, vascular amyloid deposition follows vasoconstriction in a distinct spatiotemporal pattern, progressing from smaller to larger microvessels in a sensitivity-dependent manner and exhibiting robust correlation with cognitive deterioration. This pathological sequence appears mediated through two distinct but complementary mechanisms: (1) impaired endothelial/pericyte-mediated amyloid metabolism and (2) persistent pericyte contraction augmenting vasoconstriction. Endothelial cells and pericytes critically regulate amyloid homeostasis in terminal microvasculature.^64^ Their functional impairment, indicated by diminished PDGFrβ and eNOS expression, dysregulates amyloid metabolism through increased production and reduced clearance.^64^ Amyloid aggregation subsequently enhances pericyte contractility, intensifying vasoconstriction and further compromising amyloid clearance,^65^ ultimately resulting in non-vascular amyloid deposition and cognitive impairment,^66^ as demonstrated in our investigation. Clinically, hypoperfusion precedes amyloid plaque development and cognitive decline in AD and VCI,^67^ potentially through the bidirectional interaction between microvascular constriction and vascular amyloid deposition demonstrated herein. Collectively, these findings establish vascular amyloid as a novel etiological factor in CCH pathogenesis, driving non-vascular amyloid deposition and contributing to cognitive deterioration. Future investigations examining endothelial cell and pericyte contributions to amyloid clearance are warranted, as these neurovascular unit cellular components represent promising therapeutic targets for CCH pathology.^65,68^ Understanding this vascular-to-parenchymal progression of amyloid pathology provides critical insights for developing targeted interventions that address the microvascular origins of amyloid-associated cognitive impairment.

Neuroinflammation and oxidative stress, commonly considered primary CCH drivers,^21^ appear to function as secondary mechanisms based on our findings, as their upregulation manifested exclusively during middle-to-late CCH stages. No investigations have directly established these processes as causative factors for the microvascular dysfunction characteristic of early CCH pathogenesis. Available evidence instead implicates BBB disruption and endothelial damage as initiators of CCH-associated inflammatory processes.^69^ Early endothelin-1 elevations mediate endothelial injury, subsequently promoting vascular adhesion molecule expression (ICAM1, VCAM1),^69^ consistent with our observed increases in middle and late CCH, respectively. Similarly, significant microglial and astrocytic activation emerges only during moderate CCH, following leukocyte infiltration facilitated by BBB compromise.^69–72^ BBB dysfunction has been associated with delayed (4-week post-2VO) oxidative stress in CCH,^69–72^ corroborating our SOD1 and nitrotyrosine observations. The delayed manifestation of neuroinflammation and oxidative stress biomarkers, coupled with their comparatively weak correlation with cognitive deficits, provides compelling evidence that these phenomena represent secondary pathophysiological responses within our CCH experimental timeframe. These processes likely emerge as downstream consequences of primary microvascular dysfunction, specifically endothelial damage and BBB compromise. This recharacterization of neuroinflammation and oxidative stress as secondary mechanisms may partially explain the limited efficacy of current therapeutic interventions targeting these pathways in CCH /VCI. Our findings suggest that effective therapeutic strategies should prioritize preservation of microvascular integrity and function to prevent the initiation of these secondary inflammatory and oxidative cascades, rather than attempting to suppress these processes once established.

In this investigation, CGRP emerged as a critical mediator of CCH pathology, demonstrating strongest correlations with cognitive impairment. CGRP depletion substantially contributed to CCH pathogenesis; supplementation (exogenous/endogenous) attenuated hippocampal vasoconstriction (∼90%) and vascular collapse (∼80%), addressing primary upstream microvascular dysfunction. This intervention elicited significant cognitive restoration (∼85%), markedly surpassing angiogenesis-targeted therapeutic approaches,^28,73^ which our findings identified as secondary compensatory responses rather than primary pathological mechanisms. CGRP intervention exhibited superior efficacy compared to direct Aβ42 targeting strategies, which achieved ∼40% memory enhancement and ∼50% reduction in Aβ42 concentrations,^74^ whereas CGRP supplementation facilitated ∼80-90% reduction in both vascular and parenchymal Aβ42. This efficacy differential likely stems from vasodilation’s comprehensive influence on endothelial and pericyte-mediated amyloid metabolism,^46,64^ simultaneously modulating both production and clearance pathways, rather than targeting solely β-secretase-mediated amyloid production.^74^ While exogenous CGRP supplementation conferred significant benefits, simultaneous modulation of multiple neuropeptides through endogenous approaches via DR demonstrated superior efficacy, supporting widespread vasoactive neuropeptide dysregulation as a primary CCH pathological mechanism. Both approaches normalized vascular and parenchymal endothelin-1 levels; however, endogenous CGRP upregulation induced ∼500% increase in SSTr5, a vasoactive neuropeptide receptor implicated in coagulation, compared to exogenous CGRP administration. This broader neuropeptide modulation likely underpins DR’s superior therapeutic efficacy, as trigeminal nerve activation triggers multiple neuropeptide release beyond CGRP,^37,75^ directly counteracting observed dysregulation. The comprehensive normalization of multiple dysregulated neuropeptide pathways appears to facilitate more complete restoration of neurovascular homeostasis than single-peptide interventions. Our findings indicate that: (1) widespread vasoactive neuropeptide dysregulation constitutes a primary driver of CCH pathology and represents a promising therapeutic target, and (2) multi-neuropeptide modulation elicits synergistic effects, yielding enhanced therapeutic outcomes compared to single-target approaches. These results highlight the potential of neuropeptide-based interventions as a novel therapeutic strategy for VCI and suggest that comprehensive restoration of neurovascular signaling may be necessary for optimal cognitive recovery.

Several limitations merit consideration. First, memory assessment relied on NOR and Y-maze tests, which, despite optimization for our longitudinal evaluation protocol, may inadequately capture the complete spectrum of cognitive impairment; implementation of additional behavioral paradigms such as Morris water maze would provide a more comprehensive cognitive assessment. Second, while our investigation examined 13 neuropeptides previously implicated in CCH pathomechanisms, our analysis could not encompass all potentially relevant neuropeptide biomarkers. Future investigations employing larger cohorts and comprehensive peptidomic approaches are necessary to extend and validate these findings.

## 5. CONCLUSION

Our data demonstrate that early, progressive vasoactive neuropeptide dysregulation drives microvascular constriction, constituting a pivotal mechanism in microvascular dysfunction and subsequent cognitive deterioration in chronic cerebral hypoperfusion. Microvascular constriction and vascular amyloid aggregation function synergistically, generating widespread microvascular dysfunction and non-vascular pathology through a pathological positive feedback mechanism culminating in cognitive impairment. The substantial cognitive improvements observed following CGRP supplementation underscore the therapeutic potential of this intervention strategy and warrant further investigation.

## Supporting information

Whole Blot Images

## ACKNOWLEDGEMENTS

This work is supported in part by the United States Army Medical Research Acquisition Activity (USAMRAA) under award# HT9425-24-1-1017, the US Army Medical Research and Materiel Command (USAMRMC) under award # W81XWH-18-1-0773, the Zoll Foundation Award, the merit-based career enhancement award at the Feinstein Institutes for Medical Research, and career development award from Advancing Women in Science and Medicine at the Feinstein Institutes for Medical Research.

## CONSENT STATEMENT

A consent statement was not necessary.

---

**Supplementary Figure 1.**
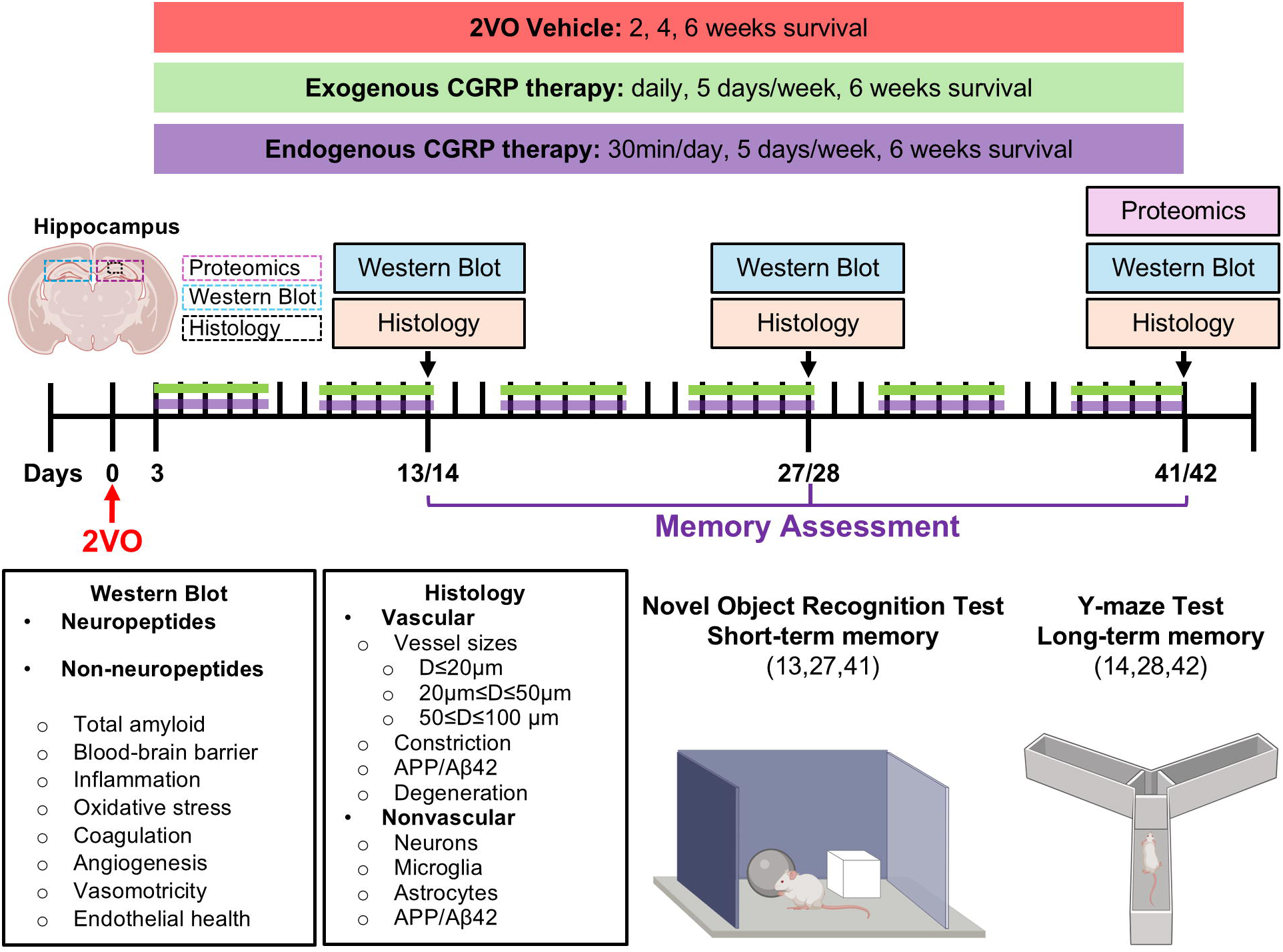
Experimental design and methodological timeline. Schematic representation of the experimental protocol timelines for vehicle (no treatment) subjects and CGRP-supplemented experimental groups. CGRP augmentation was achieved via two distinct methodologies: (1) exogenous administration utilizing intranasal delivery, and (2) endogenous upregulation through diving reflex stimulation. The protocol depicts the temporal sequence for assessment of cognitive parameters and pathophysiological endpoints. Neurocognitive function was quantified using validated behavioral assays, specifically novel object recognition and Y-maze spontaneous alternation paradigms. Pathological evaluation comprised comprehensive analysis of multiple parameters, including neuropeptide biomarkers, non-peptidergic biomarkers, microvascular structural integrity, and parenchymal damage indicators. Analytical methodologies employed a complementary multi-platform approach integrating proteomic analysis, western immunoblotting, and quantitative immunofluorescence microscopy techniques. (2VO: bilateral common carotid artery occlusion; Aβ42: amyloid β42; APP: amyloid precursor protein; CGRP: calcitonin gene-related peptide; D: diameter; \**p*□<□0.05, \*\**p*□<□0.01, \*\*\**p*□<□0.001, \*\*\*\**p*□<□0.0001)

**Supplementary Figure 2.**
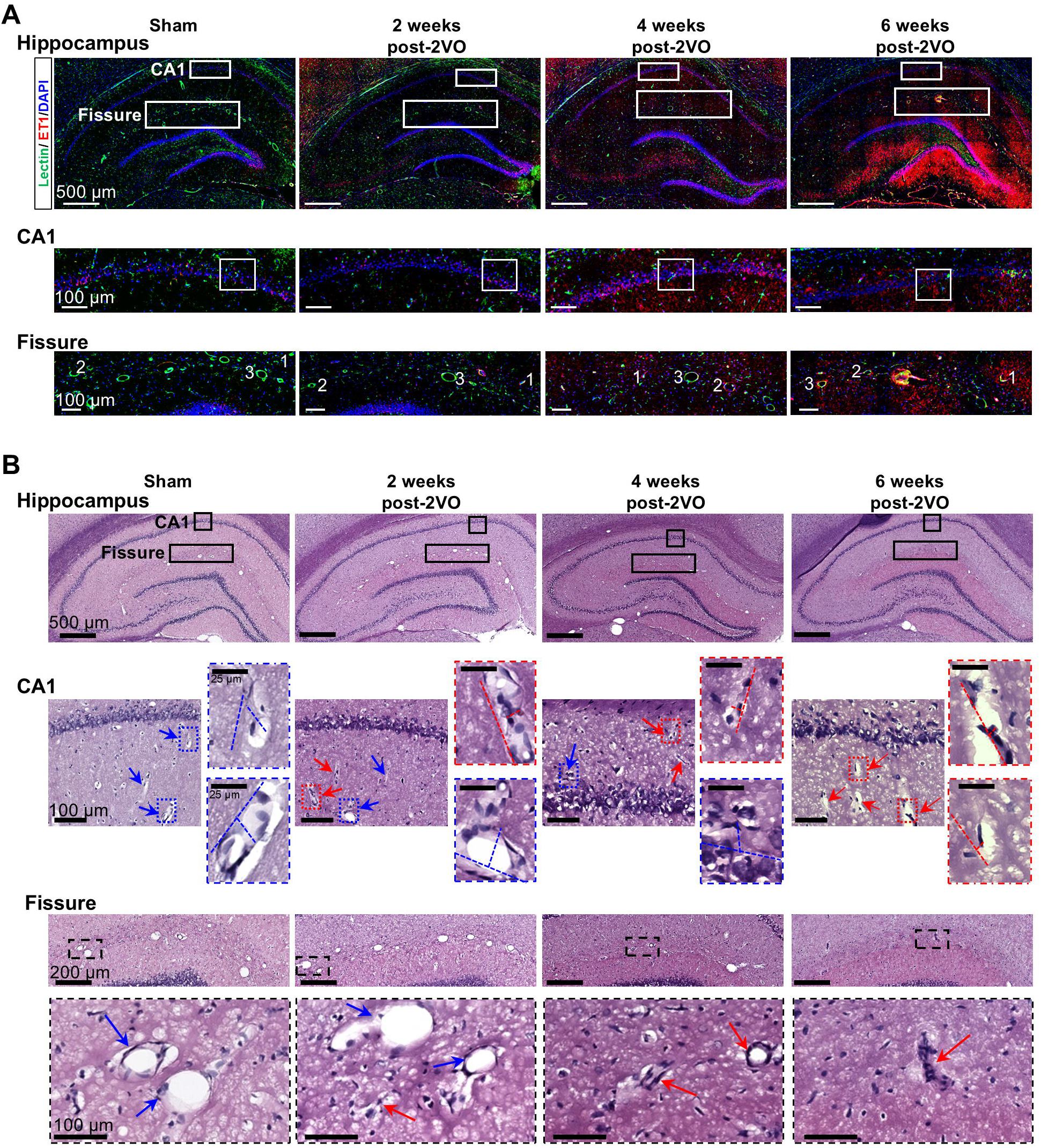
Progressive microvascular deterioration in the hippocampus following chronic cerebral hypoperfusion. Longitudinal assessment of microvascular structural alterations and vasoconstriction in the CA1 region and hippocampal fissure of rats subjected to 2VO at 2, 4, and 6 weeks post-surgery, as evaluated by immunofluorescence microscopy and histological examination. (A) Representative immunofluorescence micrographs demonstrating temporal progression of ET-1 expression co-localized with lectin-labeled microvasculature in the CA1 and hippocampal fissure, indicating progressive accumulation of this vasoconstrictor peptide. (B) Hematoxylin and eosin (H&E) stained sections illustrating time-dependent microvascular collapse and constriction within the CA1 subfield and hippocampal fissure microvasculature following CCH. (arrow = collapsed vessel, red = collapsed vessel, blue = open vessel, dotted lines = vessel width and length) (CCH: chronic cerebral hypoperfusion; 2VO: bilateral common carotid artery occlusion; ET-1: endothelin 1; DAPI: 4’,6-diamidino-2-phenylindole; H&E: hematoxylin and eosin; \**p*□<□0.05, \*\**p*□<□0.01, \*\*\**p*□<□0.001, \*\*\*\**p*□<□0.0001)

**Supplementary Figure 3.**
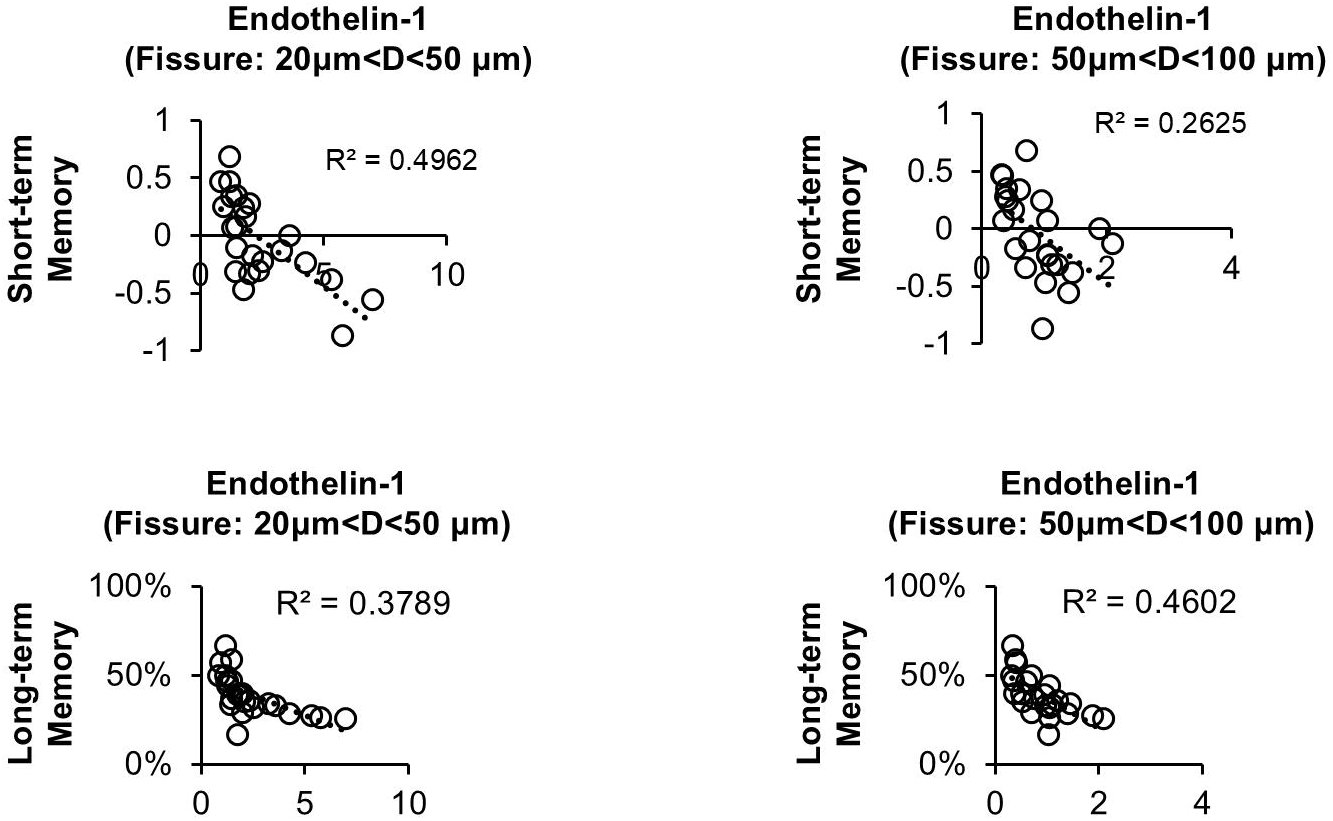
Correlation between microvascular constriction and cognitive deterioration in chronic cerebral hypoperfusion. Regression analyses examining the relationship between microvascular endothelin-1 immunoreactivity and cognitive performance measures in the chronic cerebral hypoperfusion model. Quantitative assessment revealed a statistically significant fair-to-moderate negative correlation between endothelin-1 expression and memory function parameters. The strength of this correlation was contingent upon microvascular diameter, with the correlation coefficient demonstrating vessel caliber-dependent variation, suggesting differential susceptibility of microvasculature to endothelin-mediated vasoconstriction as a potential mechanism underlying cognitive impairment.

**Supplementary Figure 4.**
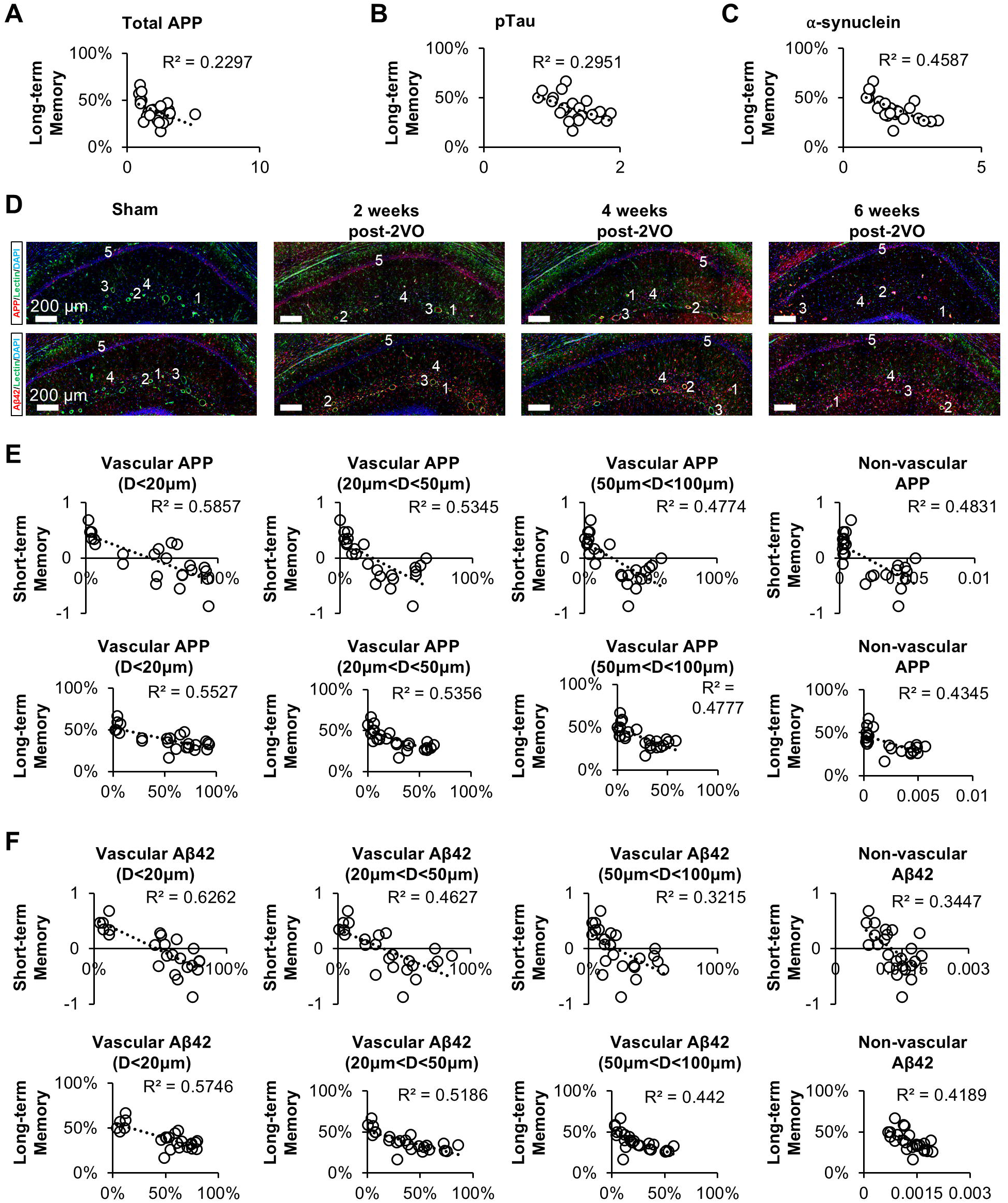
Differential correlation of vascular-associated versus total amyloidogenic protein expression with cognitive impairment in chronic cerebral hypoperfusion. **(A-C)** Regression analyses examining the relationship between total hippocampal amyloidogenic protein burden and long-term spatial memory performance in the chronic cerebral hypoperfusion (CCH) model. Quantification of total amyloid precursor protein (APP), phosphorylated tau (pTau), and α-synuclein immunoreactivity demonstrated not strong correlations with cognitive performance metrics. **(D)** Representative immunohistochemical visualization of APP and Aβ42 distribution in hippocampal tissue following CCH induction (numerical indicators correspond to panels in Figure 4D, 4E of the main manuscript). **(E-F)** Comparative correlation analyses of vascular-localized versus parenchymal APP expression with vessel diameter parameters and cognitive assessment scores. Correlation coefficients exhibited vessel caliber-dependency, with significant negative correlations observed in small-diameter microvessels and progressively diminishing correlation strength with increasing vessel diameter, culminating in non-significant associations in non-vascular parenchymal regions. These findings suggest compartment-specific contributions of amyloidogenic protein accumulation to cognitive dysfunction following CCH. (CCH: chronic cerebral hypoperfusion; APP: amyloid precursor protein; pTau: phosphorylated tau, Aβ42: amyloid β42; ET-1: endothelin 1; \**p*□<□0.05, \*\**p*□<□0.01, \*\*\**p*□<□0.001, \*\*\*\**p*□<□0.0001)

**Supplementary Figure 5.**
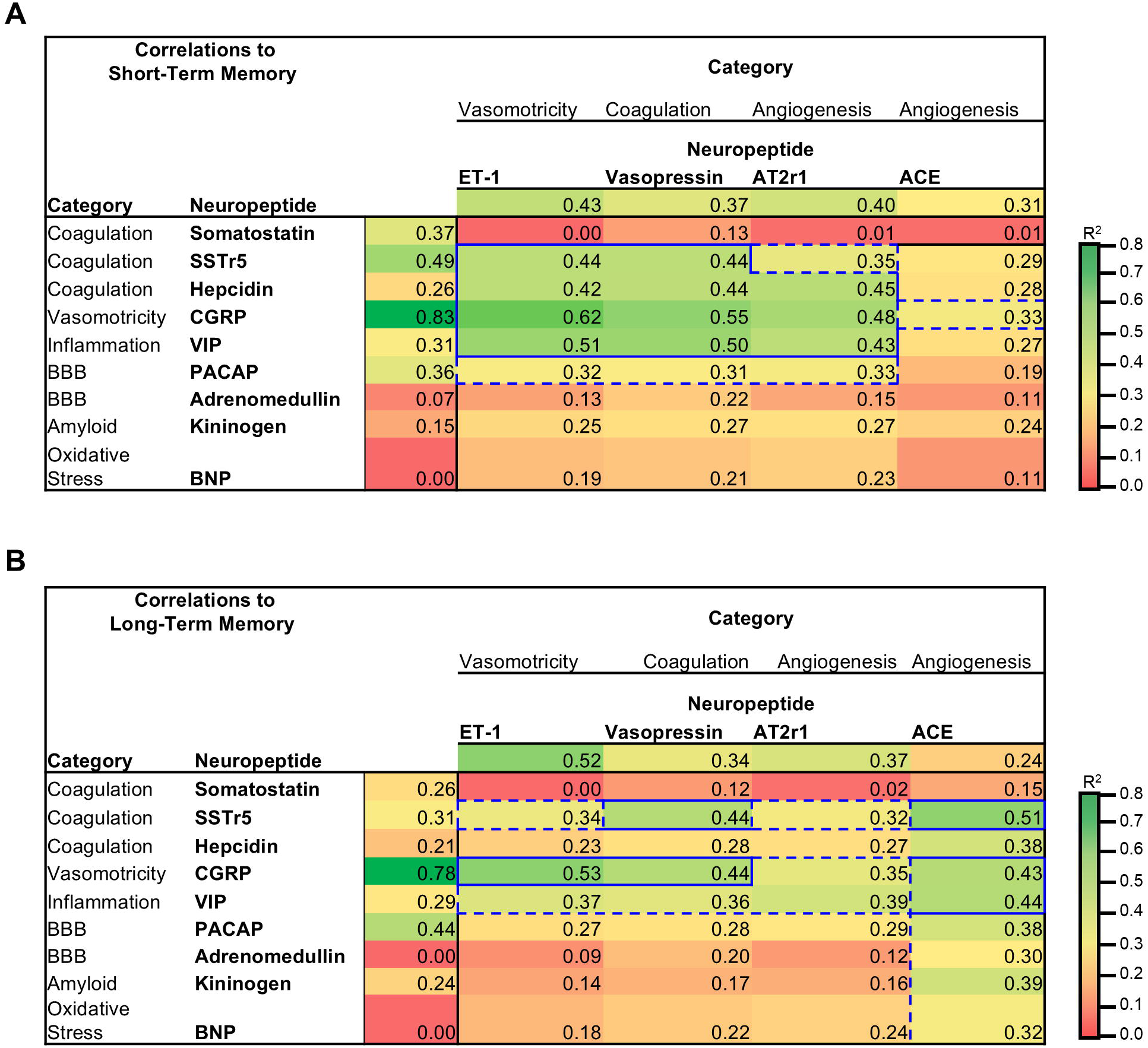
Vasomotor neuropeptide dysregulation demonstrates superior correlation with progressive memory deficits in comparison to other neuropeptide categories. Comprehensive correlation analyses examining the relationships between various neuropeptide expression profiles and cognitive performance metrics in CCH model. **(A)** Correlation matrices between absolute neuropeptide concentrations or neuropeptide expression ratios and short-term memory performance parameters. Statistical analysis revealed that predominantly vasoactive neuropeptides exhibited the strongest correlations with short-term memory impairment, with calcitonin gene-related peptide (CGRP) demonstrating the most robust association. **(B)** Correlation analyses between neuropeptide expression (both absolute values and inter-peptide ratios) and long-term memory function. Quantitative assessment indicated that primarily vasomotor regulatory neuropeptides, particularly CGRP, maintained the strongest relationship with long-term memory performance, although the correlation pattern exhibited greater variability compared to short-term memory parameters. These findings suggest differential temporal sensitivity of memory domains to vasomotor neuropeptide dysregulation following CCH. (CCH: chronic cerebral hypoperfusion; 2VO: bilateral common carotid artery occlusion; HEP: hepcidin; SST: somatostatin; SSTr5: somatostatin receptor 5; ET-1: endothelin-1; CGRP: calcitonin gene-related peptide; Vaso: vasopressin, AT2r1: angiotensin II receptor 1; ACE: angiotensin converting enzyme; VIP: vasoactive intestinal peptide; PACAP: pituitary adenylate cyclic-activating polypeptide; ADR: adrenomedullin; \**p*□<□0.05, \*\**p*□<□0.01, \*\*\**p*□<□0.001, \*\*\*\**p*□<□0.0001)

**Supplementary Figure 6.**
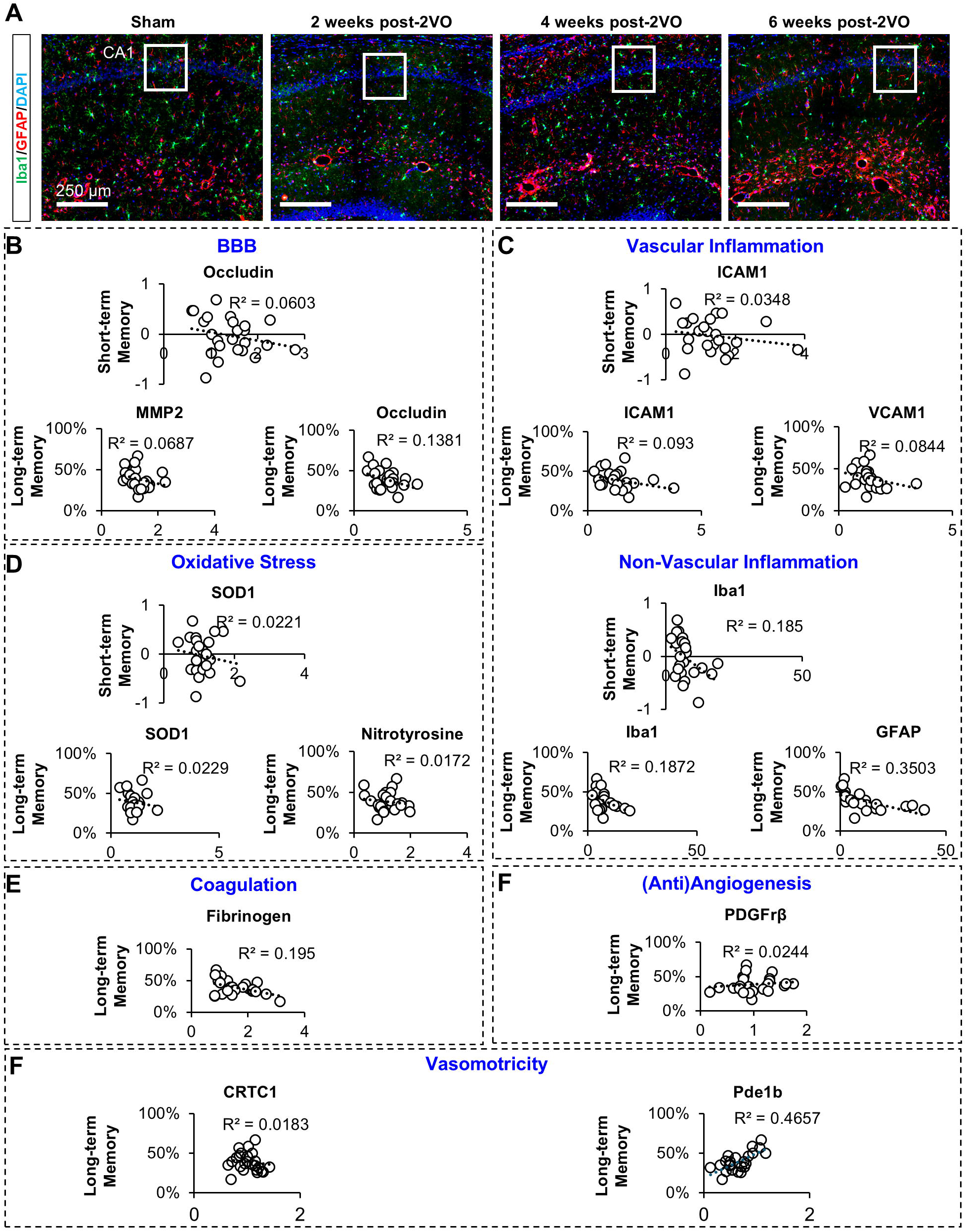
Minimal correlation between nonvascular pathological features and cognitive dysfunction following chronic cerebral hypoperfusion. **(A)** Representative immunofluorescence micrographs illustrating reactive gliosis in the hippocampal formation, with visualization of activated astrocytes and microglia following induction of CCH. **(B-F)** Correlation analyses examining relationships between quantified nonvascular pathological indicators and cognitive performance parameters. It revealed predominantly weak correlations across multiple markers of parenchymal injury. These findings suggest that, unlike vascular pathology, nonvascular tissue alterations contribute minimally to the cognitive impairment observed in this experimental model of CCH. (CCH: chronic cerebral hypoperfusion; 2VO: bilateral common carotid artery occlusion; MMP2: matrix metalloproteinase 2, BBB: blood brain barrier; ICAM1: intercellular adhesion molecule 1; VCAM1: vascular cell adhesion molecule 1; SOD: superoxide dismutase; NT: nitrotyrosine; PDGFrβ: platelet derived growth factor receptor β; Pde1b: Phosphodiesterase 1B; CRTC1: CREB-regulated transcription coactivator 1; Iba1: ionized calcium binding adaptor molecule 1; GFAP: glial fibrillary acidic protein)

**Supplementary Figure 7.**
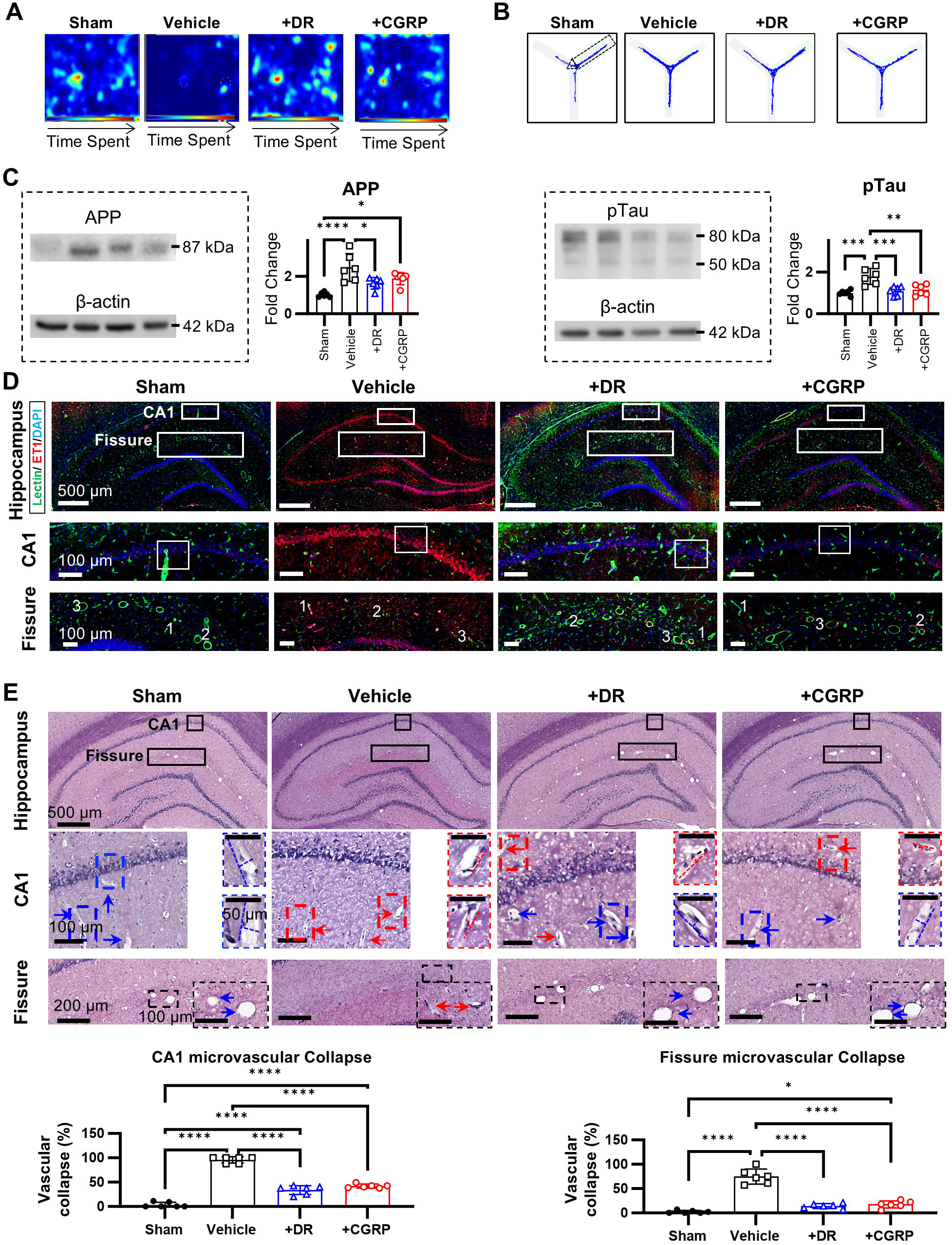
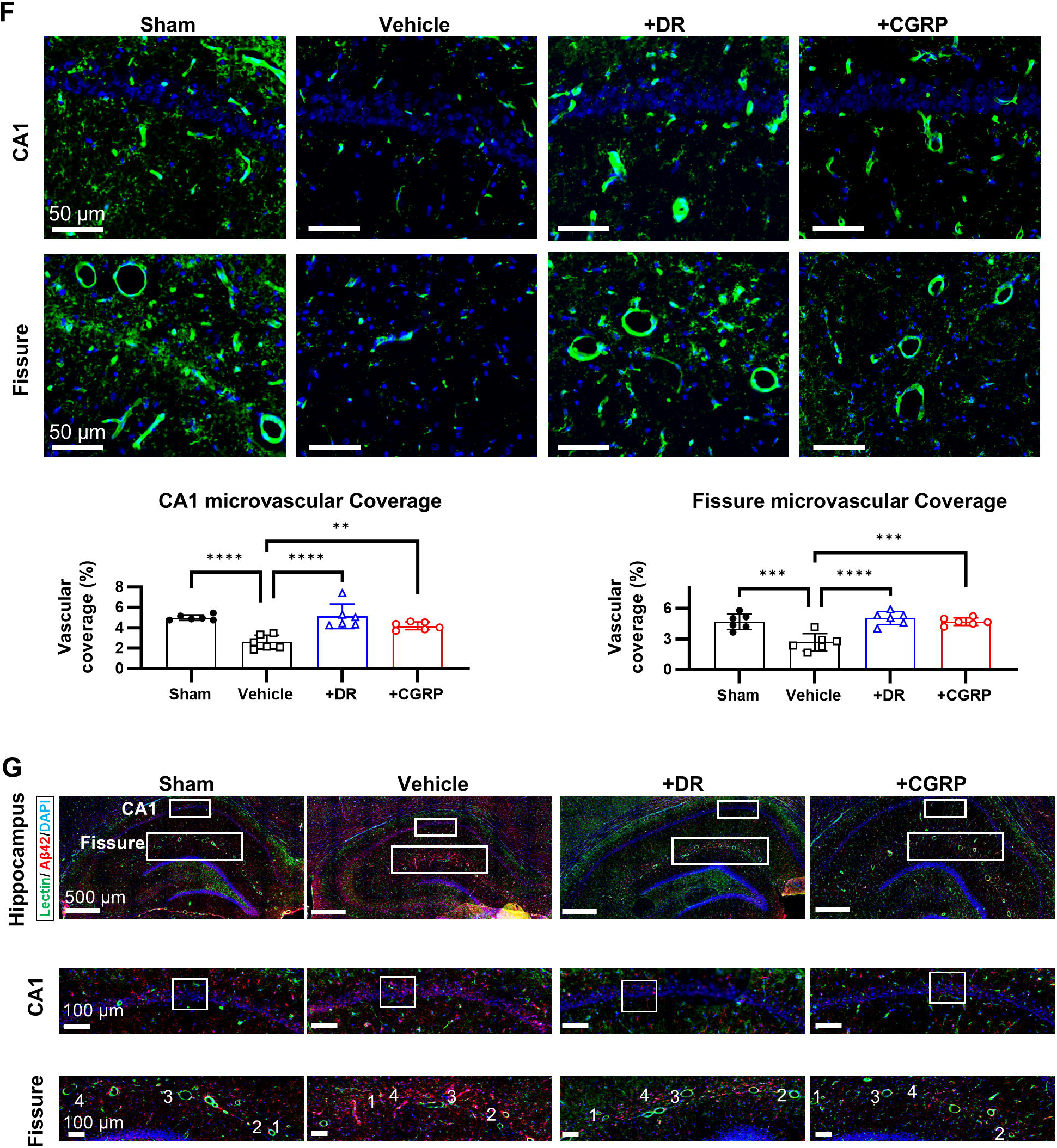
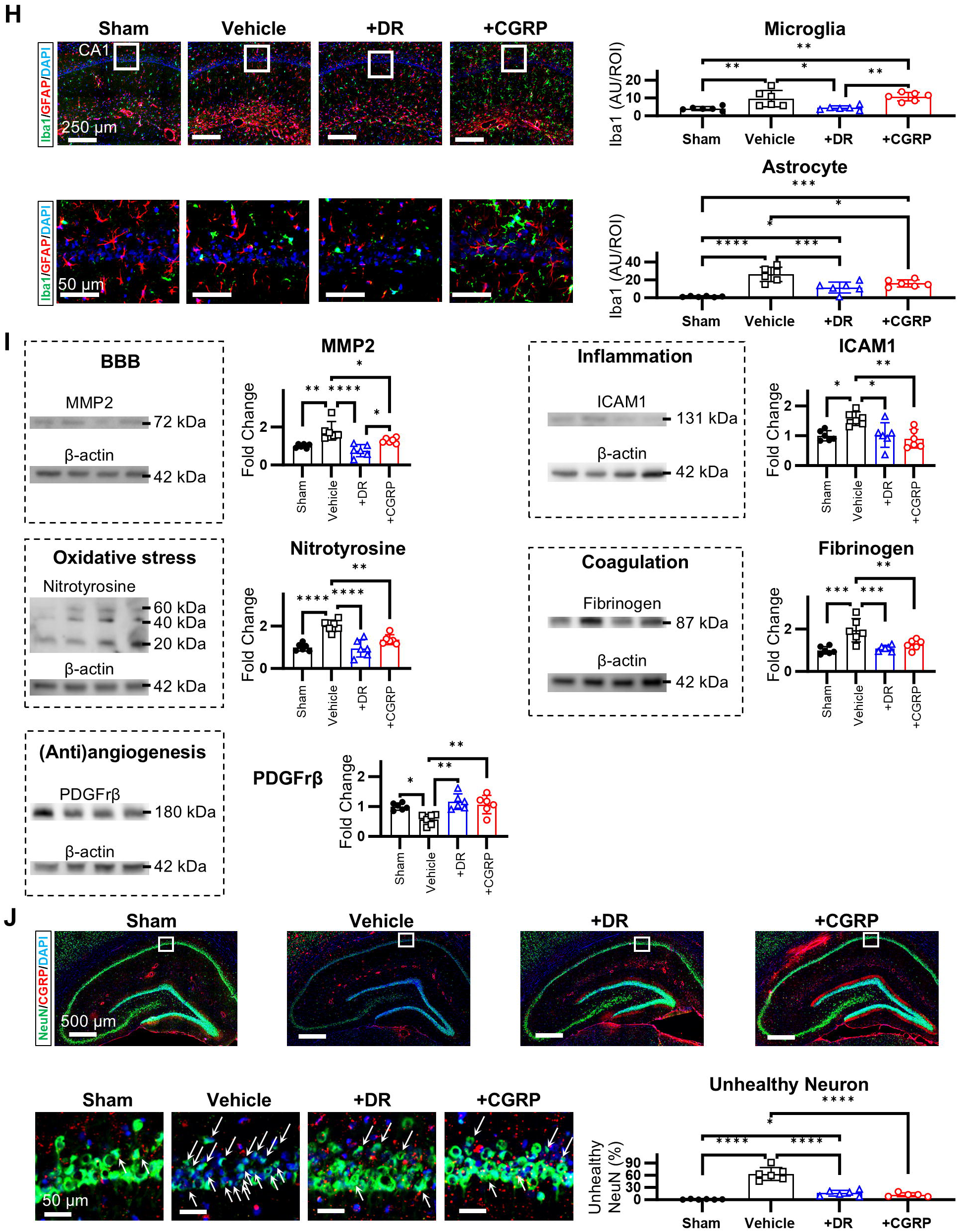
CGRP Supplementation Mitigates Cognitive Dysfunction Through Attenuation of Microvascular and Non-vascular Pathologies. **(A-B)** CGRP supplementation significantly ameliorated both short-term and long-term memory deficits at 6 weeks following bilateral common carotid artery occlusion (2VO). **(C)** Densitometric analysis demonstrated that endogenous and exogenous CGRP modulation significantly downregulated amyloid precursor protein (APP) and phosphorylated Tau (pTau) expression at 6 weeks post-2VO. **(D)** Immunofluorescence imaging revealed that both endogenous and exogenous CGRP supplementation attenuated endothelin-1 (ET-1) expression at 6 weeks post-2VO. **(E)** Semi-quantitative analysis demonstrated that CGRP supplementation significantly mitigated microvascular collapse within the CA1 and fissure regions of the hippocampus. **(F)** Quantitative immunofluorescence analysis showed that endogenous and exogenous CGRP supplementation significantly increased microvascular coverage. **(G)** Immunofluorescence imaging demonstrated that CGRP administration decreased APP and Aβ42 expression throughout the hippocampus. **(H)** Quantitative immunofluorescence analysis revealed that endogenous and exogenous CGRP supplementation decreased both Iba1 and GFAP expression at 6 weeks post-2VO, indicating reduced neuroinflammation. **(I)** Densitometric analysis indicated that CGRP modulation mediated the amelioration of both vascular and non-vascular pathogenesis. **(J)** Quantitative immunofluorescence assessment demonstrated that CGRP supplementation significantly improved neuronal viability in the CA1 subregion of the hippocampus. (CCH: chronic cerebral hypoperfusion; 2VO: bilateral common carotid artery occlusion; MMP2: matrix metalloproteinase 2, BBB: blood brain barrier; ICAM1: intercellular adhesion molecule 1; VCAM1: vascular cell adhesion molecule 1; SOD: superoxide dismutase; NT: nitrotyrosine; PDGFrβ: platelet derived growth factor receptor β; Pde1b: Phosphodiesterase 1B; CRTC1: CREB-regulated transcription coactivator 1; Iba1: ionized calcium binding adaptor molecule 1; GFAP: glial fibrillary acidic protein; \**p*□<□0.05, \*\**p*□<□0.01, \*\*\**p*□<□0.001, \*\*\*\**p*□<□0.0001) (arrow = collapsed vessel, red = collapsed vessel, blue = open vessel, dotted lines = vessel width and length)

