## Supplementary material for "Vasoactive Neuropeptide Dysregulation: A Novel Mechanism of Microvascular Dysfunction in Vascular Cognitive Impairment": Whole Blot Images

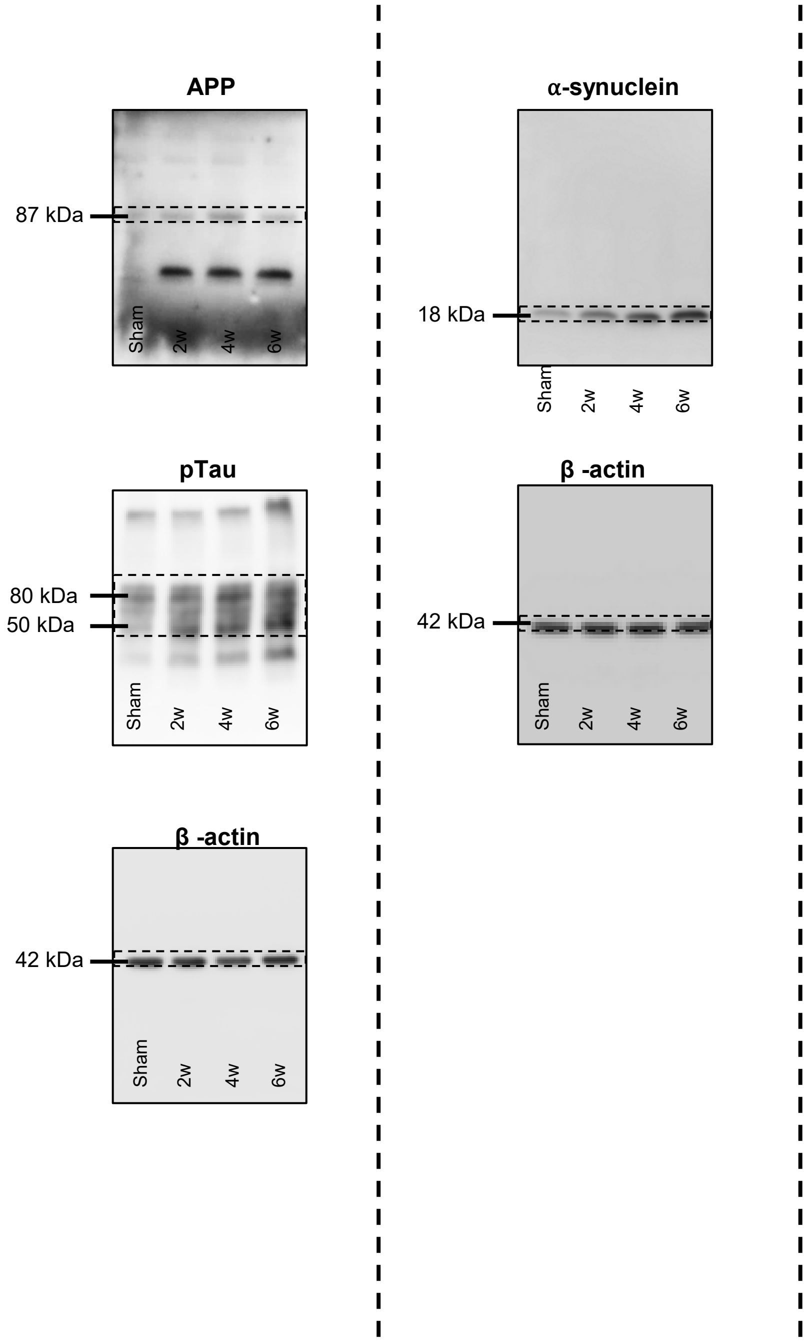


### Figure 4 – Whole Blot Images

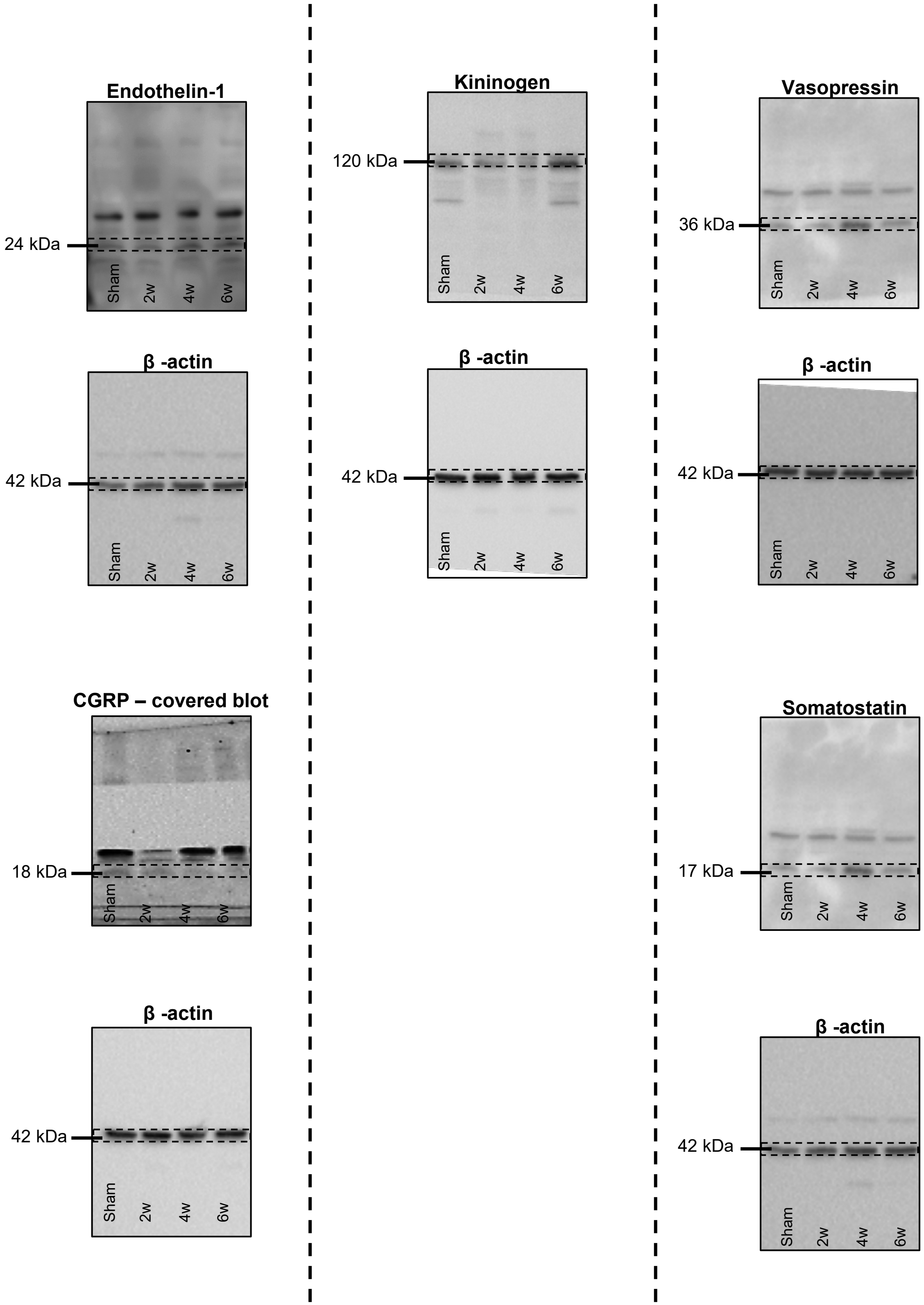


### Figure 5 – Whole Blot Images, P1

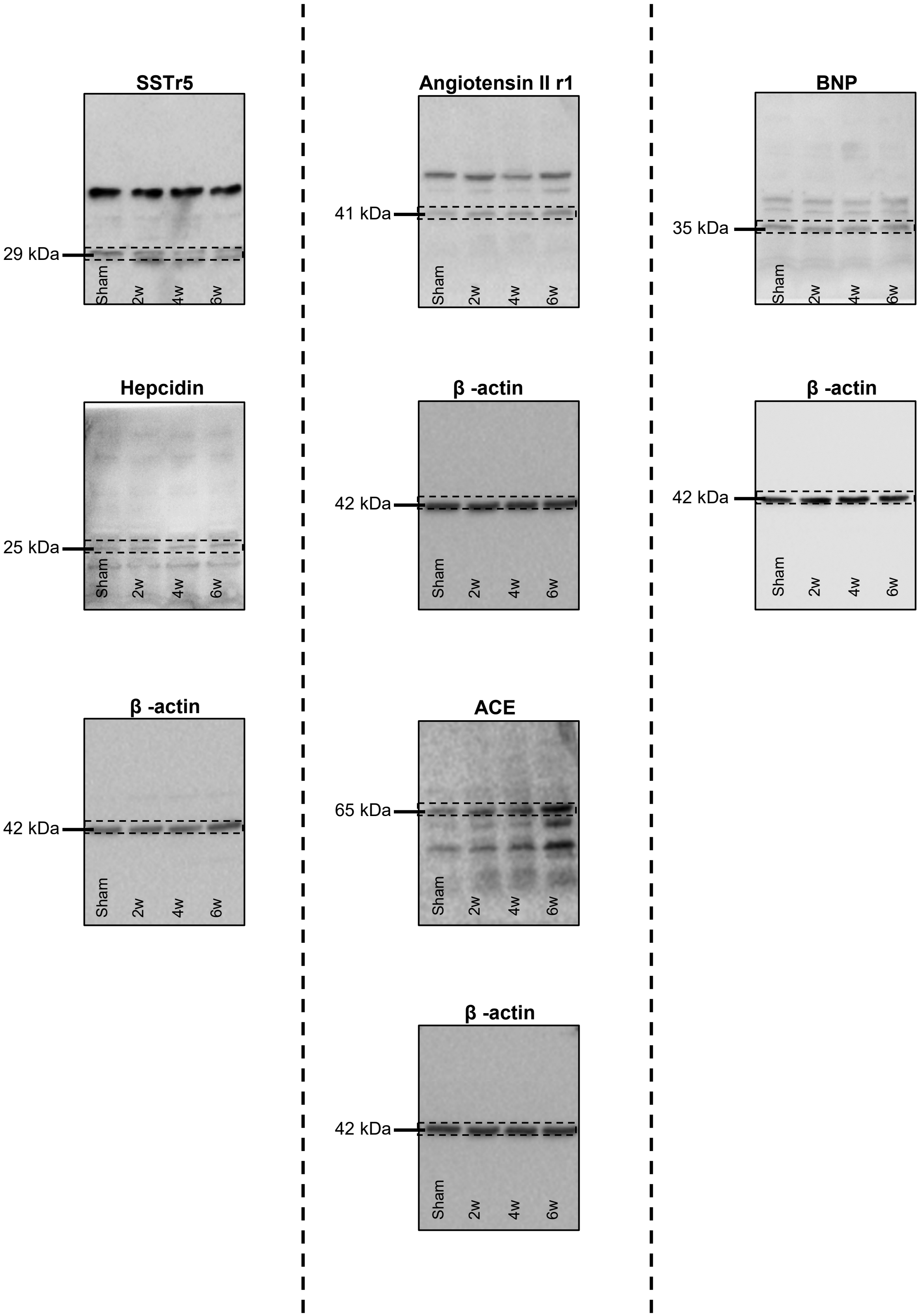


### Figure 5 – Whole Blot Images, P2

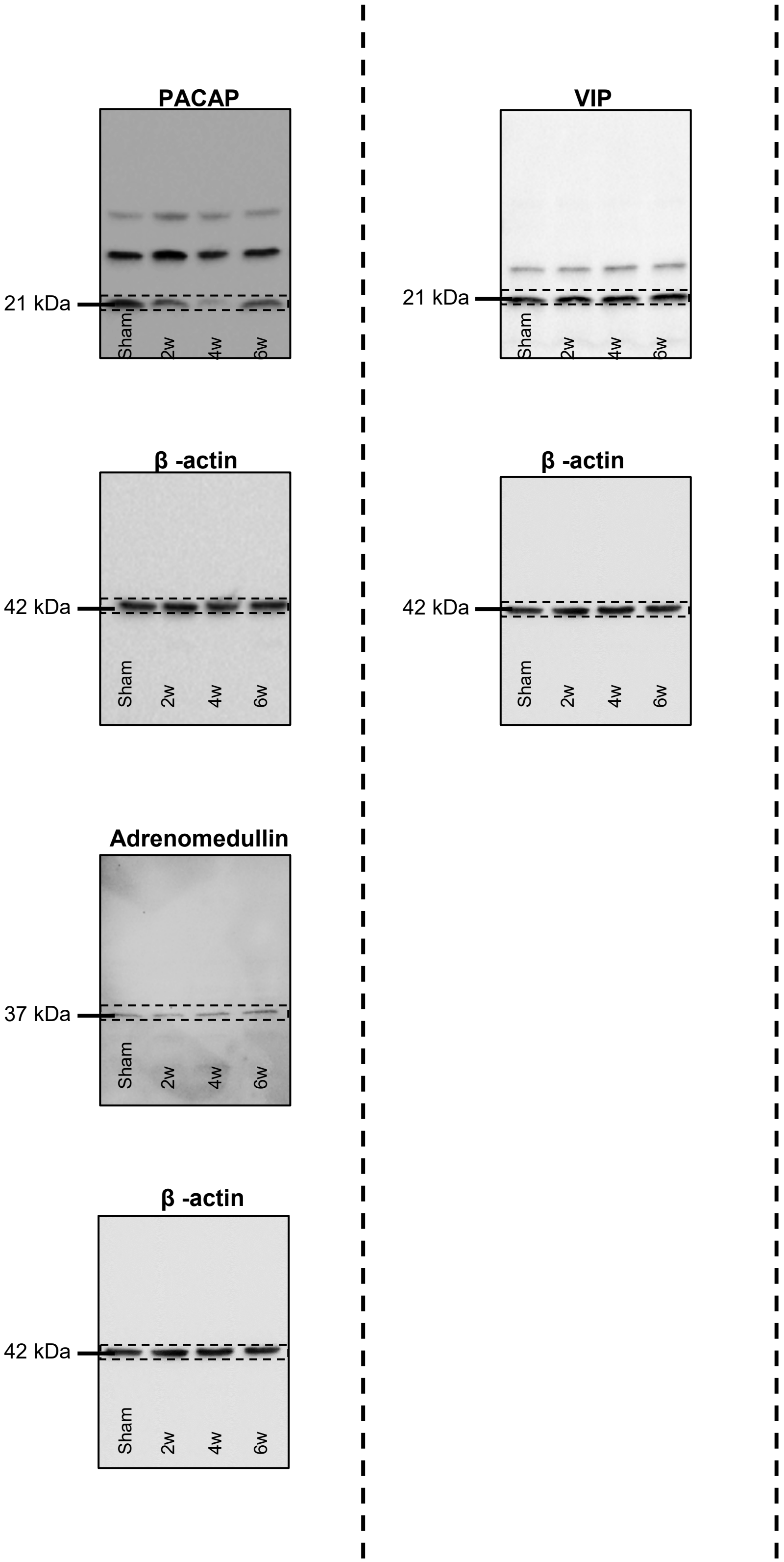


### Figure 5 – Whole Blot Images, P3

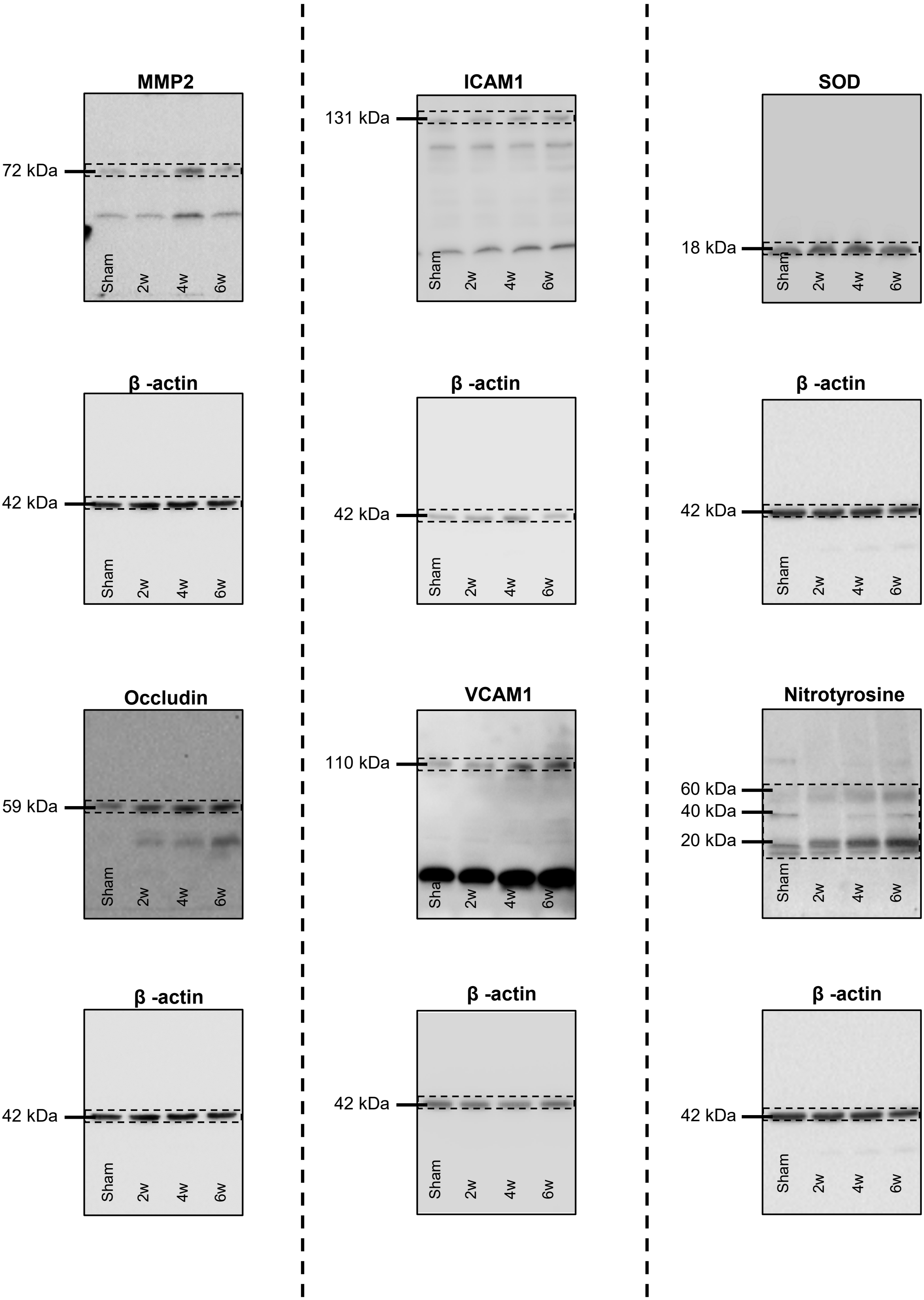


### Main Figure 6 – Whole Blot Images, P1

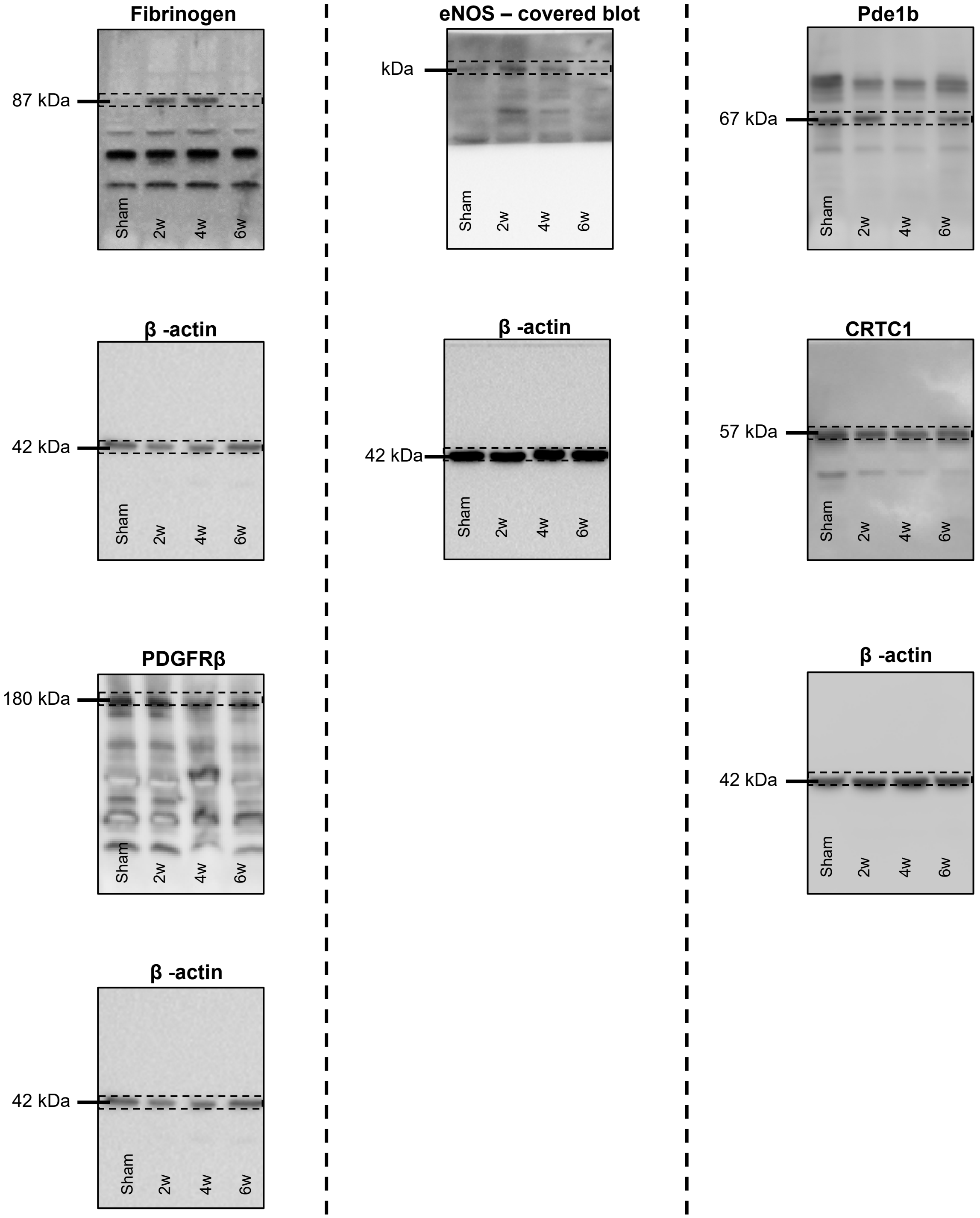


### Main Figure 6 – Whole Blot Images, P2

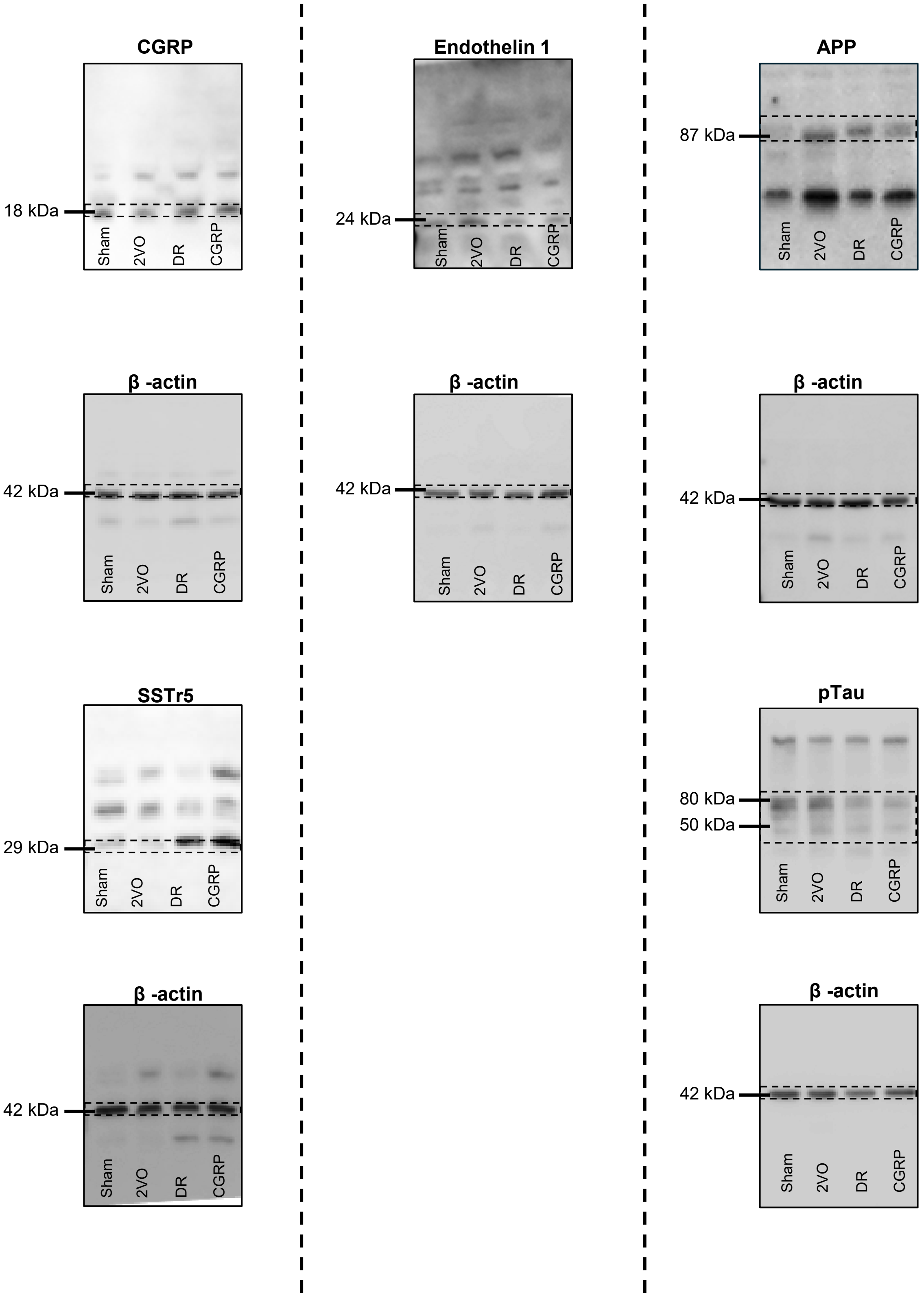


### Main Figure 8 – Whole Blot Images, Part 1

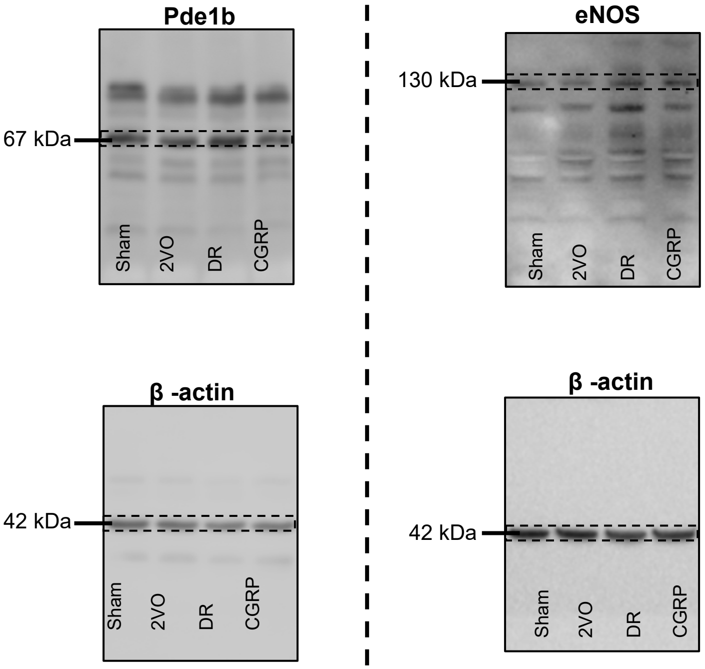


### Main Figure 8– Whole Blot Images, Part 2

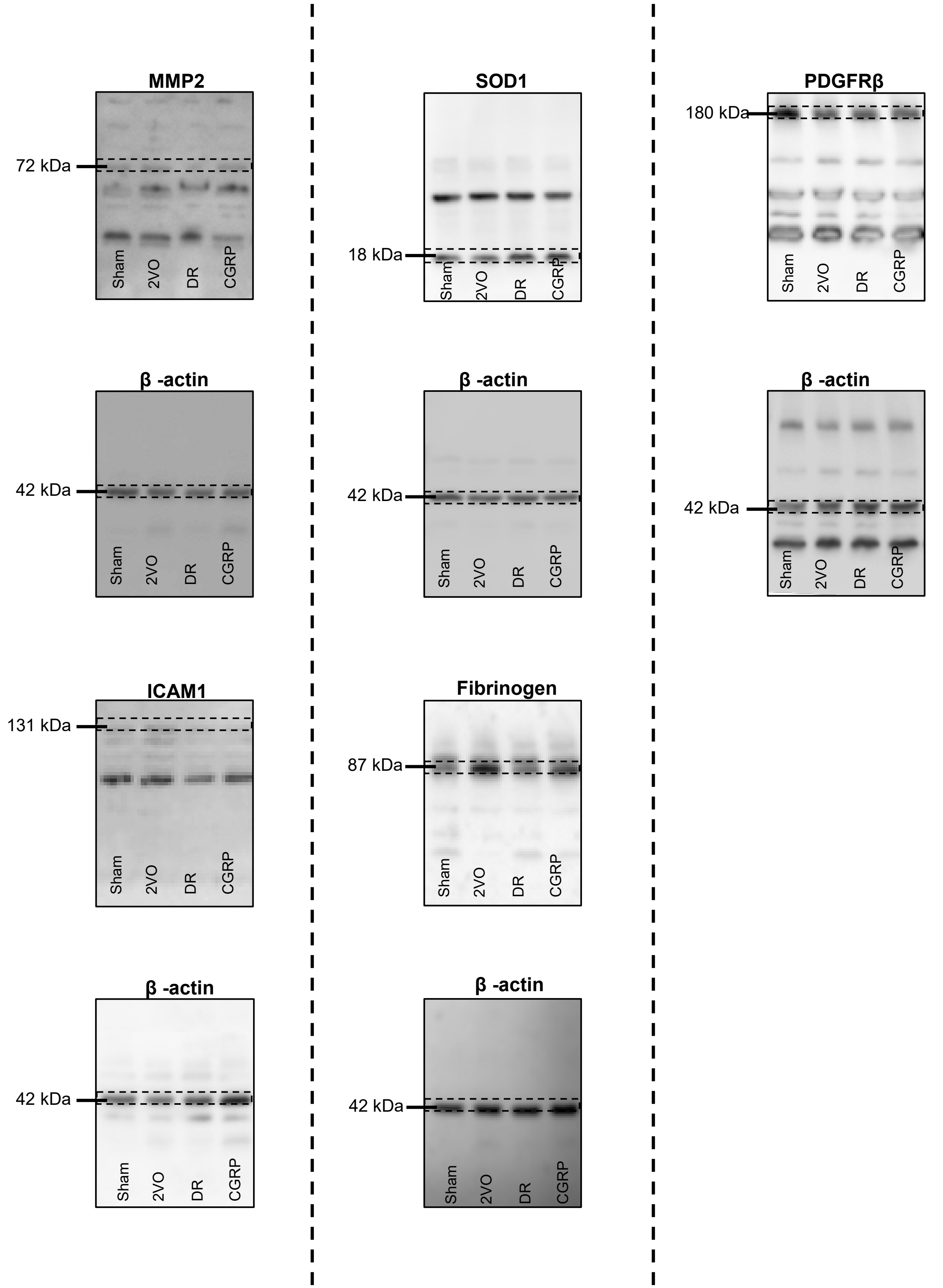


### Supplemental for Main Figure 8 – Whole Blot Image
