## Supplemental Tables for "Vasoactive Neuropeptide Dysregulation: A Novel Mechanism of Microvascular Dysfunction in Vascular Cognitive Impairment"

**Table 1**. Antibodies used for western blot analysis.

| **Antibody** | **Host** | **Dilution factor** | **Function(s)**  *(Primary; secondary; tertiary)* | **Catalog#** | **Company** |
| --- | --- | --- | --- | --- | --- |
| ACE | Rabbit | 1:1000 | Vascular degeneration; Inflammation; Vasomotricity | 24743-1-AP | Proteintech, USA |
| ADR | Rabbit | 1:1000 | BBB; Vasomotricity; Angiogenesis | AB190819 | Abcam, USA |
| ADR 2 | Rabbit | 1:1000 | BBB; Vasomotricity; Angiogenesis | 23781-1-AP | Proteintech, USA |
| AT2r1 | Rabbit | 1:1000 | Vascular degeneration; Inflammation; Vasomotricity | 25343-1-AP | Proteintech, USA |
| BNP | Rabbit | 1:1000 | Oxidative stress; Vasomotricity | AB19645 | Abcam, USA |
| CGRP | Mouse | 1:100 | Vasomotricity; Angiogenesis;  Synaptic Plasticity; Neurotransmission | Sc-374221 | Santa Cruz BioTechnology, USA |
| ET-1 | Rabbit | 1:1000 | Coagulation; Vasomotricity; Synaptic Plasticity | 12191-1-AP | Proteintech, USA |
| Fibrin(ogen) | Mouse | 1:500 | Coagulation | AB119948 | Abcam, USA |
| Hepcidin | Rabbit | 1:1000 | Coagulation; BBB; Vasomotricity | AB30760 | Abcam, USA |
| ICAM1 | Rabbit | 1:1000 | Vascular Inflammation | 10831-1-AP | Proteintech, USA |
| Kininogen | Rabbit | 1:1000 | Amyloid-dependent inflammation; Vasomotricity | 11926-1-AP | Proteintech, USA |
| MMP2 | Mouse | 1:1000 | BBB | 66366-1-Ig | Proteintech, USA |
| Nitrotyrosine | Mouse | 1:1000 | Oxidative Stress | AB61392 | Abcam, USA |
| Occludin | Mouse | 1:1000 | BBB | 66378-1-Ig | Proteintech, USA |
| PACAP | Rabbit | 1:5000 | BBB; Vasomotricity; Angiogenesis | AB181205 | Abcam, USA |
| PDGFrβ | Rabbit | 1:1000 | Angiogenesis | 13449-1-AP | Proteintech, USA |
| SOD1 | Rabbit | 1:2500 | Antioxidant | 10269-1-AP | Proteintech, USA |
| SST | Rabbit | 1:1000 | Coagulation; Vasomotricity;  BBB | AB307803 | Abcam, USA |
| SSTr5 | Mouse | 1:1000 | Coagulation; Vasomotricity;  BBB | 66772-1-ig | Proteintech, USA |
| Vasopressin | Rabbit | 1:1000 | Vasomotricity; BBB;  Neurotransmission | AB213708 | Abcam, USA |
| VIP | Rabbit | 1:1000 | Inflammation; Vasomotricity; Angiogenesis | 162333-1-AP | Proteintech, USA |
| APP | Mouse | 1:1000 | Amyloid formation | Ab126649 | Abcam, USA |
| pTau | Rabbit | 1:1000 | Stabilizes neuronal microtubule | AB32057 | Abcam, USA |
| α-synuclein | Rabbit | 1:1000 | Control of neurotransmitter release | 2642S | Cell Signaling Technology |
| VCAM1 | Rabbit | 1:1000 | Vascular inflammation | 30958-1-AP | Proteintech, USA |
| Pde1b | Rabbit | 1:1000 | Modulates intracellular cGMP and cAMP (Vasomotricity) | 13121-1-AP | Proteintech, USA |
| CRTC1 | Rabbit | 1:1000 | CREB regulator (Vasomotricity) | 10441-1-AP | Proteintech, USA |
| eNOS | Rabbit | 1:1000 | Endothelial Cell Health | PA1-037 | Invitrogen, USA |
| β-actin | Mouse | 1:25000 | N/A | A5441 | Sigma, USA |
| α-Mouse Secondary, HRP | Goat | 1:2000 | N/A | Ab6789 | Abcam, USA |
| α-Rabbit Secondary, HRP | Goat | 1:1000 | N/A | Ab97051 | Abcam, USA |

* ACE: angiotensin converting enzyme, ADR: Adrenomedullin, APP: amyloid precursor protein, AT1: angiotensin II receptor 1, BNP: brain natriuretic peptide, cAMP: cyclic adenosine monophosphate, cGMP: cyclic guanosine monophosphate, CGRP: calcitonin gene-related peptide, CREB: cAMP response element binding protein, CRTC1: CREB-regulated transcription coactivator 1, eNOS: endothelial nitric oxide synthase, ET-1: endothelin 1, PACAP: pituitary adenylate cyclase activating peptide, Pde1b: phosphodiesterase 1B, pTau: phosphorylated tau, SST: somatostatin, SSTr5: somatostatin receptor 5, VIP: vasoactive intestinal peptide,

**Table 2**. Antibodies used for immunofluorescent analysis.

| **Antibody** | **Host** | **Dilution factor** | **Catalog #** | **Company** |
| --- | --- | --- | --- | --- |
| Lectin | N/A | 1:75 | FL-1171-1 | Vector Laboratories, USA |
| ET-1 | Rabbit | 1:1000 | 12191-1-AP | Proteintech, USA |
| DAPI | N/A | 1:2000 | 62248 | Thermo Fisher Scientific, USA |
| GFAP | Mouse | 1:100 | ab4648 | Novus Biologicals, USA |
| Iba1 | Rabbit | 1:400 | GTX100042 | Genetex, USA |
| APP | Mouse | 1:200 | ab126649 | Abcam, USA |
| Aβ42 | Rabbit | 1:200 | Bs-0107r | Bioss, USA |
| pTau | Rabbit | 1:100 | ab32057 | Abcam, USA |
| NeuN | Mouse | 1:500 | ab104224 | Abcam, USA |
| CGRP | Mouse | 1:50 | Sc-374221 | Santa Cruz BioTechnology, USA |

Aβ42: amyloid β 42, APP: amyloid precursor protein, CGRP: calcitonin gene-related, DAPI: 4',6-diamidino-2-phenylindole, ET-1: endothelin 1, GFAP: glial fibrillary acidic protein, Iba1: calcium-binding protein, pTau: phosphorylated Tau protein, protein
